# Reconstructing the network of horizontal gene exchange in bacteria to differentiate direct and indirect transfers

**DOI:** 10.1101/2025.11.20.689518

**Authors:** Michael Sheinman, Tommaso Stentella, Paul Etheimer, Florian Massip, Peter F. Arndt

## Abstract

Horizontal gene transfer (HGT) plays a central role in bacterial evolution. Yet, its large-scale dynamics and underlying network structure remain poorly characterized. We present a theoretical framework that models HGT as a continuous stochastic process over a network of bacterial genera and analyze its genomic footprint via the distribution of exact sequence matches shared across taxa—the match length distribution (MLD). We show that different evolutionary regimes imprint distinct statistical signatures on the MLD: single episodic gene transfer events yield exponential distributions, while continuous sustained HGT processes lead to power-law tails. The power-law exponent is analytically linked to the topology of the transfer network, distinguishing between intraclade transfers and hub-mediated dissemination. Empirical MLDs derived from bacterial genomes recapitulate these predicted patterns. Moreover, we find that defining a genus-specific “transferability” parameter that governs pairwise HGT rates and incorporating a high-transferability hub accurately reproduces the observed data. Our approach provides a general framework for inferring hidden structure in genomic horizontal transfer processes, enabling quantitative analysis of microbial evolution.

## I. INTRODUCTION

The most influential concepts in biological evolution— such as the phylogenetic tree [1] and the molecular clock [2]—are not readily applicable to the bacterial domain of life. In bacteria, evolutionary relationships are often better described by a *phylogenetic web* rather than by a strictly bifurcating tree [3, 4], and molecular divergence fails to consistently correlate with speciation time [5]. These deviations from canonical models are largely attributed to the pervasive influence of *horizontal gene transfer* (HGT) [6].

The widespread occurrence of HGT has reshaped our understanding of microbial evolution, giving rise to the concept of a web of life [3, 7]. While the existence of HGT is well established, its large-scale organization and the statistical signatures it leaves in genome sequences remain poorly understood. It was found that some bacterial lineages function as phylogenetic hubs, receiving and donating genes more extensively than others [8]. However, the relative importance of such transfer hubs—and, more generally, of indirect transfer paths compared to direct exchanges—has never been characterized [9]. Previous studies have used different measures of genomic similarities between different bacteria to identify and quantify homologous recombination and gene transfer [10– 13]. Such measures of shared homologous sequences can then be used to build weighted HGT networks (see *e*.*g*. [14–20]). In these and other studies, usually, when the shared homology cannot be explained by conservation, an edge between the nodes is established. Using such an approach, direct and indirect transfers cannot be distinguished. Furthermore, edges represent a similarity measure but not the direct HGT rate.

In the present study we analyze maximal exact sequence matches shared by genomes to infer the HGT network. Long exact matches have been observed between highly divergent bacterial taxa, such as *Escherichia* and *Salmonella* [21–23]. Since their presence cannot be explained by vertical inheritance under typical bacterial mutation rates [24–27], they are the remnant of past HGT. Such long matches are easy to find and they provide a strong signal of recent gene transfer [28–32]. However, the presence of a long exact match between two distant genomes does not necessarily imply a direct transfer between the two *i* →*j*. It could have been mediated by a third taxon (*i* →*k* →*j*) or acquired independently from a shared donor (*i* ← *k* →*j*), as shown in Fig. 1. These possibilities naturally suggest viewing HGT as a *network-level process*, where gene flow occurs both directly and indirectly through interconnected taxa. To disentangle direct and indirect transfers, we develop a quantitative model of HGT as a *continuous stochastic process on a genus-level network*. In our model, edges represent the direct HGT rate, in contrast to a measure of similarity. Note that bacteria of a genus might live in different environments [33]; nevertheless, our model considers only the network of bacterial taxa (genera) and does not differentiate between bacteria within a genus.

**FIG. 1:**
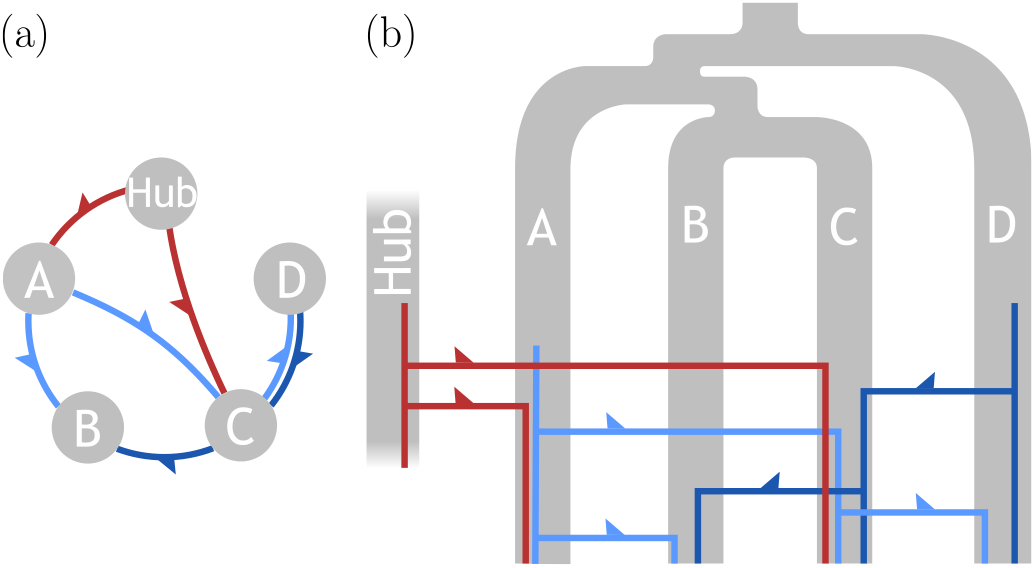
Illustration of (a) HGT dissemination scenarios between four distant taxa, *A, B, C* and *D*, and an un-sampled “hub” taxon, resulting in (b) three phylogenetic trees (red and blue lines). Thick grey lines represent the speciation history of the taxa. We assume that all the speciation events occurred long before the recent HGT events which dominate the long exact matches between the genomes of the taxa. The HGTs are shown as arrows in (a). The total (vertical, to the most recent common ancestor) length of each HGT resulting tree in (b) is defined as *τ*.

To explore this idea, we analyze the *match length distributions* (MLDs) of maximal exact sequence matches shared among sets of *n* ≥ 2 genera. Extending previous work [30, 32], we show that the shape of the MLD encodes key information about the structure of the underlying HGT process. A single transfer event produces an exponential distribution; multiple discrete events generate mixtures of exponentials; and a continuous HGT process yields a *power-law tail* of the match length distribution, *m*(*r*) ~ *r*^−*α*^. Crucially, the exponent *α* reflects the topology of the transfer network. Notably, it is sufficient to represent a bacterial genus as a node with a characteristic *transferability* parameter, such that the HGT rate between any two genera is given by the product of their transferabilities. This simple, minimal model leads to concrete, testable predictions about the statistical properties of exact sequence matches shared between genomes.

In the following sections, we first quantify the extent of multi-genera exact matches (Section II). We then derive analytical predictions for match-length distributions under a continuous HGT model and show how indirect transfers via a hub modify the scaling exponents (Section III). Then we fit the model to all observed MLDs (Section IV). Finally, we discuss mechanistic interpretation and broader implications (Section V).

## II. IDENTIFYING HGT EVENTS AFFECTING MULTIPLE GENERA

To investigate the exchange of genetic material, we began by identifying exact sequence matches—DNA segments that are identical across genomes from multiple genera within the *Enterobacteriaceae* family and the *Serratia* genus. Following [30], we restricted our analysis to very long matches (≥ 1000 bp): such a high threshold was chosen to ensure we exclude matches resulting from evolutionary conservation [32]. We first detected exact matches shared between pairs of genomes and then extended this analysis to identify regions shared among larger sets of genera (*i*.*e*., sequences identical in *n* genera, with *n* ranging from 3 to 8).

Under neutral expectations (if matches shared between 2 genera A and B were independent from the matches shared between 2 other genera C and D), the number of shared matches should decline rapidly as the number of compared genera increases. Surprisingly, we observed a substantial number of long exact matches shared among three or more species. For example, on average, about 5% of the *Escherichia* genome participates in long exact matches shared with *any* one of the eight genera considered, while 0.39% of the *Escherichia* genome is shared by all eight genera considered (i.e., present in at least one genome in each of the eight genera), whereas under the independence-of-matches hypothesis this number should be much lower (of the order of 5%^7^ ≃ 10^−9^ « 0.39%). Other genera have similar fractions (see Supplemental Figure S3). As shown in Supplemental Figure S4, genes located within these long shared regions are enriched in mobile genetic elements [34]. This confirms that those matches are the result of HGT events and not due to contamination or artifacts of the method.

To gain further insight into the HGT processes, we calculated the distribution of the lengths of matches shared by different sets of genera (see Methods for details). In [30] we showed that the analysis of the MLD obtained by comparing pair of bacterial species allowed to measure their exchange rate. Thus we tested whether we could apply the same methodology to matches shared by a larger set of species. To our surprise, the MLDs constructed from matches shared by more than two species followed power-law distributions with exponent different from the *α* = −3 values obtained in our first study (see Fig. 3(a)). Indeed, the exponent of the power law decreases with the number of species considered (see Fig. 3(b)).

In the following, we propose a simple model that explains these empirical observations and demonstrate that we can use this model to disentangle the contributions of direct and indirect transfers and measure direct and indirect transfer exchange rates between the eight genera considered in our study.

## III. THE MODEL AND ITS ANALYSIS

In this section we present the model and derive analytical results for the expected MLD for a given HGT network. Before doing so, it is instructive to analyze a single HGT event spreading in a set of taxa *s* and the shape of its MLD.

### A. A single HGT event

Consider a mobile DNA sequence of a certain length *L* that spreads through the set *s* of taxa. Any dissemination scenario (see Fig. 1(a)) of such mobile element can be presented as a phylogenetic tree with *n* = |*s*| leaves (see Fig. 1(b)). Defining the total length of this tree by *τ* « 1 and assuming a constant mutation rate, defined as 1 in our time units, the probability that a bp along the DNA segment is identical for all set members is *e*^−*τ*^. Then, the expected number of maximal exact matches of length *r « L* decays exponentially with *r* [35]:

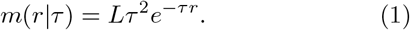

Therefore, for a single HGT that has disseminated in all the studied genera, we expect the length distribution of the exact matches shared by all genomes to be exponential. As a consequence, when several HGT events occur, they should generate an MLD distributed as a mixture of several exponential distributions whose parameter depends on *τ*.

Assuming that HGT is a frequent process, we expect that *τ* has a certain continuous density *P*_*s*_(*τ*) for a given HGT network. Then, the MLD of this set is given by

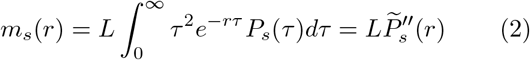

where 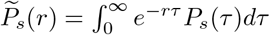 is the Laplace transform of *P*_*s*_(*τ*). One can see that the length distribution of exact sequence matches is related to the Laplace transform of the distribution of the total length of the phylogenetic trees of the HGT events. Hence, studying the MLD allows us to evaluate the density *P*_*s*_(*τ*). In the following, we show that *P*_*s*_(*τ*) informs us on the structure of the HGT network. We then study empirical observations of *m*_*s*_(*r*), to validate the model and fit its parameters for considered genera.

### B. Continuous HGT model

The central insight is that exact-match lengths *r* reflect the distribution of total divergence times *τ*, which in turn depends on the HGT network. By modeling HGT as a continuous-time process over a weighted network, we can derive 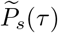and thus the MLD *m*_*s*_(*r*) analytically. To do so we model HGT via a weighted network where nodes are bacterial taxa (we use genera in this article, but in this section we keep it more general). A pair of taxa that exchange DNA via HGT are connected by an edge whose weight represents the effective rate (per bp) of HGT between the taxa pair. Effective rates take into account that the transferred gene is not present in 100% of the genomes of the donor taxon and is not transferred to 100% of genomes of the recipient taxon. We assume that all transfers are Poisson processes with different rates. Our analysis focuses on very long exact matches (> 1000 bp) shared by distant taxa. Hence, the contribution of conservation to the distribution we study is negligible. Throughout the paper we define time units so that the mutation rate *µ* = 1, and all inferred effective HGT rates are given in units of a typical bacterial mutation rate.

We consider a network of taxa exchanging genetic material that belongs to a mobilome (a collection of all mobile elements) [36, 37] of a given length. The exchange rate per bp from taxon *i* to taxon *j* is denoted by *ρ*_*ij*_ and is assumed to be the same for all elements composing the mobilome. It is possible to compute the analytical solution for a general asymmetric transfer rate matrix (see Eqs. (S5,S13) in the Supplemental Results S.1), but the solution is then difficult to fit to empirical data. We assume that every genus *i* is characterized by its transferability *γ*_*i*_, such that the transfer matrix is symmetric and rank-1, given by *ρ*_*ij*_ = *γ*_*i*_*γ*_*j*_. Interestingly, this simpler, special case is sufficient to fit the empirical data. We find that it is useful to obtain asymptotic results in the large *r* limit. For the considered system the asymptotic formula is very close to the exact one but has the advantage of being easier to interpret.

#### 1. Asymptotic results

For any connected set *s*, in the asymptotic regime of large *r* the MLD of the set depends only on the transfer rates within the set (see Supplemental Results, Section S.1.4). The explicit closed form solution of the MLD 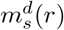 (*d* for transfers only within the set *s*) can be obtained as shown in Section S.1.4: in a *s*-set of HGT with |*s*| = *n* taxa with transfer rates *ρ*_*ij*_ the MLDs are given by

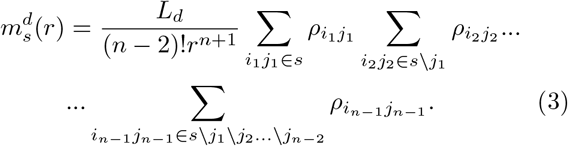

One can see that in the asymptotic regime the MLD of the set *s* depends only on the HGT rates within the set *s. L*_*d*_ is the size of the mobilome that is exchanged directly between the considered genera. The scaling law for the long-match asymptotics is

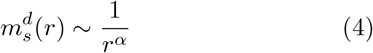

where *α* = *n* + 1 = |*s*| + 1. For pairwise comparisons, *n* = 2, one gets the known result *α* = *n* + 1 = 3 [30]. As we show below, the prediction *α* = *n* + 1 for *n* > 2 does not match the empirical data. This discrepancy suggests that for larger genera sets *s*, HGT events are increasingly dominated by an additional, indirect HGT pathway that involves organisms outside the focal set. In the following, we show that such pathway can be explained by a common unobserved network hub.

#### 2. Adding a strong hub to the network

Consider a network hub (with index *i* = 0) that is connected to a set *s* with *n* genera, labeled as *i* = 1, …, *n*, with rates *ρ*_0*i*_ = *γ*_0_*γ*_*i*_ (see Fig. 2). In Section S.1.5 we show that the hub contribution to the MLD in the limit *r* → ∞ is given by

**FIG. 2:**
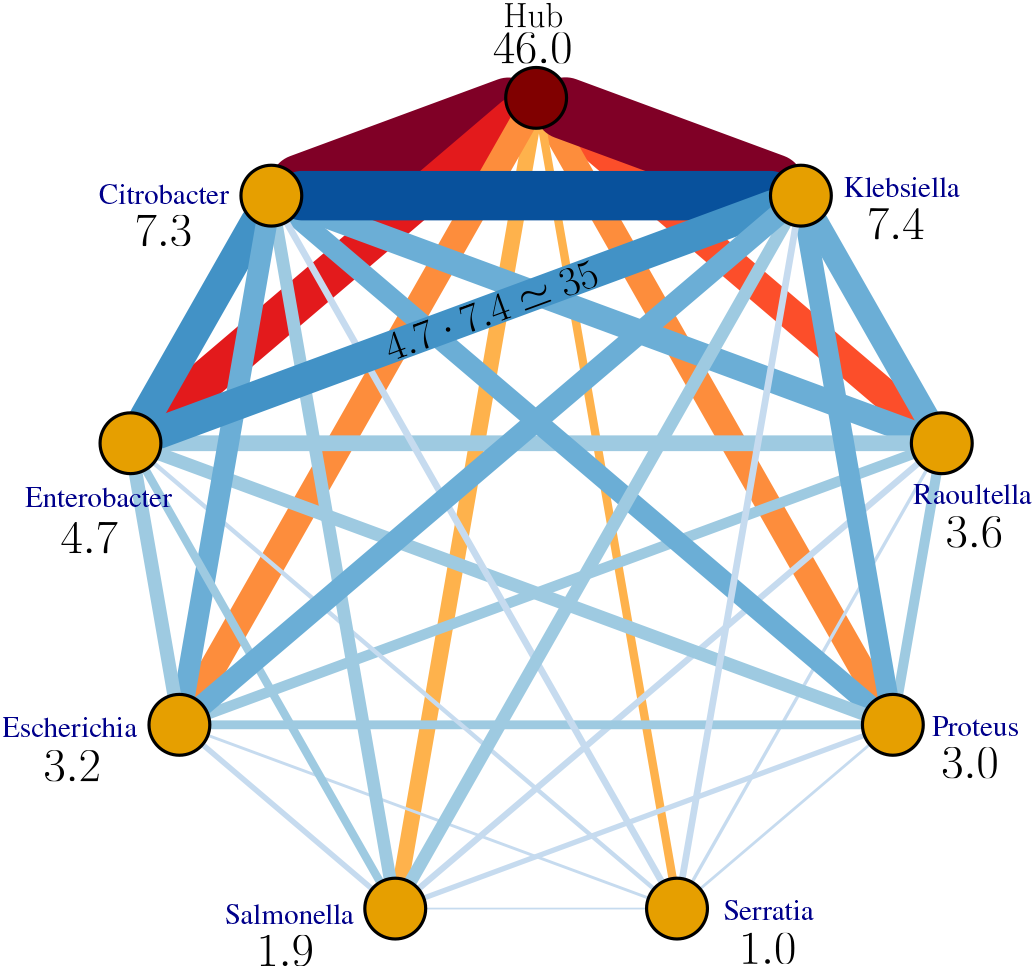
Inferred network of 8 bacteria genera. Thickness of the lines and color intensity is proportional to HGT rate *ρ*_*ij*_. The edges from the hub are presented with the shades of red and scaled down by a factor of 5 relative to other edges (presented with the shades of blue), for a better visibility. The numbers under the genera names indicate genes transferability *γ*_*i*_. The edge between *Enterobacter* and *Klebsiella* exemplifies the assumption of the model that the HGT rate between two genera is given by the product of their tranferabilities, *ρ*_*ij*_ = *γ*_*i*_ · *γ*_*j*_. In addition to the fitted values of *γ*_*i*_ presented on this plot, the fitted direct mobilome size is *L*_*d*_ *~* 10^5^ and the size of the mobilome that goes through the hub is fitted to *L*_*h*_ *~* 2.2 · 10^4^.

**FIG. 3:**
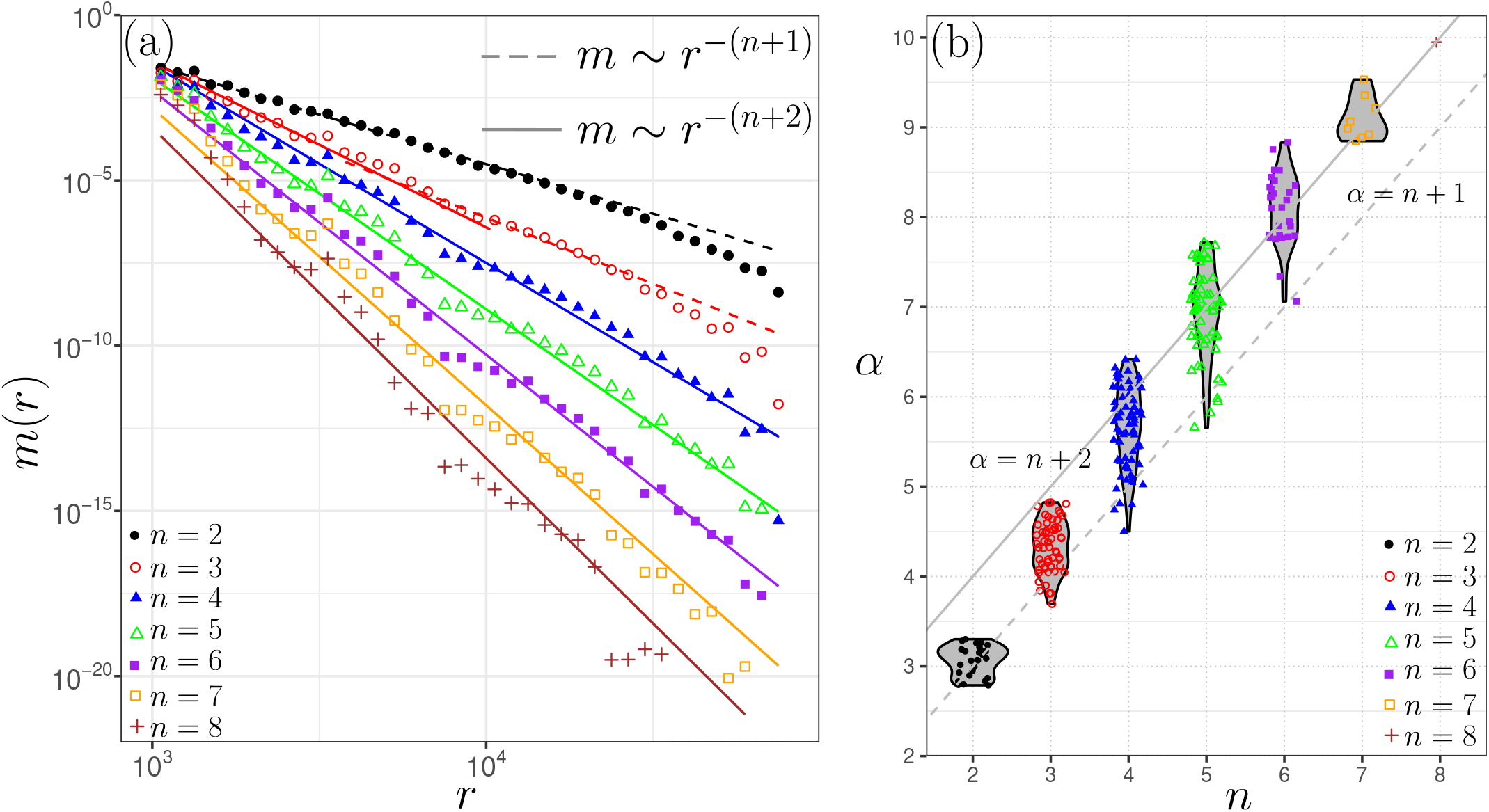
(a) Means of MLDs of all comparisons of pairs (*n* = 2), trios (*n* = 3), quartets (*n* = 4) *etc*. The dashed line represents the power-law scaling *r*^−(*n*+1)^, while the solid line represents the power-law scaling *r*^−(*n*+2)^. Green lines is fitting with sum of two (for *n* = 10) or three (for *n* = 9) exponents. The insets represent the distribution of the power-laws *α* for all comparisons with a given *n* using the method of moments. (b) Violin plot of the power-law estimates of the MLDs for all comparisons of pairs (*n* = 2), trios (*n* = 3), quartets (*n* = 4) *etc*. Dashed line represents *α* = *n* + 1 (only transfers within the set) and solid line represents *α* = *n* + 2 (transfers via a hub outside of the set).

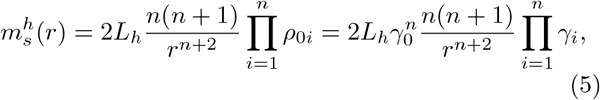

where *L*_*h*_ is the size of the mobilome that is exchanged via the hub. The scaling law of this hub-mediated long matches asymptotics is

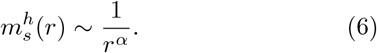

where *α* = *n* + 2 = |*s*| + 2. One can also consider the hub as a common source, such that *ρ*_0*i*_ = *γ*_0_*γ*_*i*_, but *ρ*_*i*0_ = 0 for each *i*. In this case the MLD is the same as in Eq. (5), without the factor 2. In the following, we will assume that the rate from the hub to a genus is equal to the rate in the opposite direction. However, the same fits of the empirical MLDs can be obtained with a common source (such that the rates to the hub are all zero), just replacing *L*_*h*_ by 2*L*_*h*_. In Sections S.1.5 and S.1.6 we provide a detailed calculation for MLD for these two cases.

#### 3. The complete MLD

The complete MLD is the sum of the contribution from direct transfers and from the indirect transfers that occur via the hub

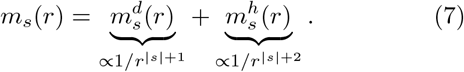

Therefore, the observed power-law exponent *α* of the empirical MLD of a given set of genera reflects the dominating HGT pathway connecting the genera in the set.

#### 4. Number of parameters of the model

For a HGT network with *n* taxa we have overall *n*(*n* − 1) rates *ρ*_*ij*_. However, since we assume that *ρ*_*ij*_ = *γ*_*i*_*γ*_*j*_ is a rank-1 symmetric matrix, the number of free parameters is the number of *n* + 1 transferabilities, *γ*_*i*_, including the hub. In addition we need to model the length of the mobilome, differentiating two set of genes: genes that are transferred directly and constitute the “direct mobilome” of size *L*_*d*_ and the set of genes that are transferred via the hub and constitute the “indirect mobilome” of size *L*_*h*_. In total we have *n* + 3 free parameby ters. The number of MLD functions that we fit is given by 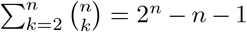. For our case of *n* = 8 genera, our model have 11 free parameters to fit 247 MLDs.

Having established the theoretical framework, we now evaluate it against empirical match-length distributions. Our model predicts that the MLD of an *n*-taxa set reflects the evolutionary history of HGT events. If HGTs are rare, the MLD will consist in a combination of several exponential distributions. If, however, HGTs are more frequent, the MLDs will be a combination of many exponentials and form a power-law:

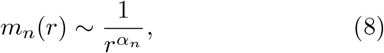

with *α*_*n*_ = *n* + 1 if most of the matches are coming from the transfers within the set and *α*_*n*_ = *n* + 2 if indirect transfers dominate. Finally, the prefactor of the powerlaw depends on the HGT rates between the genera. In this Section we test these theoretical predictions on empirical data and infer the effective HGT rates.

We determine the 11 free parameters of the model by jointly fitting all 247 considered MLDs, yielding the network shown in Fig. 2. The theoretical predictions show good agreement with the empirical distributions, supporting the validity of our framework. For *n* = 2, the predicted exponent *α* = *n* + 1 = 3 indicates that the MLDs are dominated by direct transfers (see Figs. 3(a,b), 4(a), S8(a), Fig. S9, and Ref. [30]).

For *n* > 2, however, the MLDs are largely dominated by matches exchanged through the hub. In particular, matches shared by *n* > 3 taxa are well approximated by power laws with exponent *α* = *n* + 2 (see Figs. 3, 4(c–g), S8(c–g), and Figs. S11–S14).

**FIG. 4:**
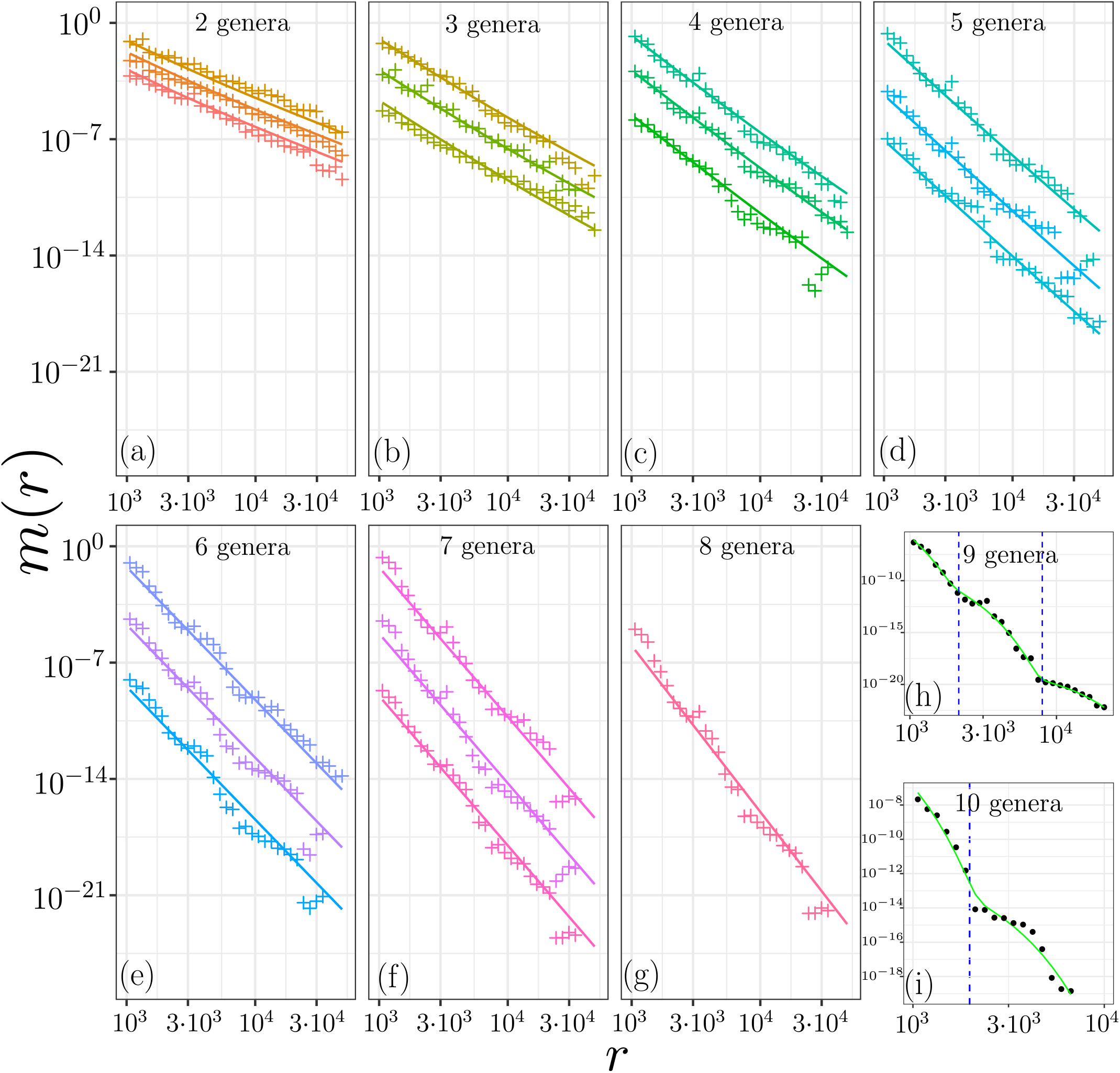
(a-g) MLDs of three selected comparisons of pairs (*n* = 2), trios (*n* = 3), quartets (*n* = 4) *etc* (see upper-right corners). Markers represent the empirical data and the lines represent the theoretical predictions. The MLDs are multiplies by constant prefactors for a better visibility, so the *y*-axes are arbitrary. For each set the empirical MLD and the theoretical one are multiplies by the same number. The number of considered sets for each *n* are shown in the lower-left corners. (h) MLD of *n* = 9 set: 8 considered genera plus *Vibrio*. The green line is fit with 3 exponential functions and the dashed lines represent the crossover between the exponential functions. (i) MLD of *n* = 10 set: all genera from (h) and *Cronobacter*. The green line is fit with 2 exponential functions and the dashed line represents the crossover between the exponential functions. All MLDs are shown in Fig. S8 and more detailed in Figs. S9,S10,S11,S12,S13,S14).

The case *n* = 3 is more nuanced. For several MLDs we observe a clear transition between two regimes (see Figs. 3, 4(c), S8(c), and Fig. S10). These distributions are well described by a sum of two power laws with exponents *α* = *n* + 1 = 4 and *α* = *n* + 2 = 5: shorter matches are dominated by within-group HGT (*α* = 4), whereas longer matches reflect indirect transfers via the hub (*α* = 5). Consequently, the estimated values of *α* for *n* = 3 scatter between 4 and 5 (see Fig. 3(b)).

### A. Matches shared by many genera (*n* = 9, 10)

Another prediction of the model is that a single HGT event generates an exponential MLD, such that rare HGT events are expected to generate an MLD that looks like a sum of a few exponents. Moreover, if each exponent corresponds to a horizontal dissemination of a single DNA segment, the matches that contribute to it are expected to form a contiguous DNA segment.

To test these predictions, we analyze the matches which are shared by *n* = 9, 10 genera, adding *Vibrio* and *Cronobacter* to the considered genera (only for this analysis). Such widespread matches appear due to HGT events that manage to spread to a significant fraction of the population in many genera. Such events are rare and, as our theory predicts, result in MLD which can be well fitted by a combination of a few exponential distributions. The MLD of matches shared by *n* = 9 genera is shown in Fig. 4(h). This MLD can be well modelled by the combination of 3 recent events of spreading of DNA segments among the strains of considered genera. On Fig. 4(i) we present the MLD of the matches shared by 10 genera. In that case the MLD is dominated by only two HGT events and can be fitted by two exponential distributions. To further support this conclusion, we analyze the sequences shared by *n* = 9, 10 genera.

We can decompose MLDs for *n* = 9 and *n* = 10 into several exponential distributions. The crossing over from one exponent to the next (blue dashed lines in Fig. 4) define length classes of matches contributing to each component. According to our theory, we expect that all matches contributing to each of those exponential components originate from the same transferred region. Thus, the DNA sequences of the matches in each component should be related.

To test this hypothesis, we assembled the matches of each of these classes (see Supp. Mat. S.2). Matches in each class overlap sufficiently so that they can be assembled into long contigs. Notably, those belonging to the class with longest mean length can be assembled into a single contig. We show that such contigs have different functional annotations. Overall, those results are in good agreement with our theory, confirming the models’ assumptions.

### B. HGT Events Are Predominantly Plasmid-Mediated

Horizontal gene transfer in bacteria typically occurs through one of three canonical mechanisms: conjugation, transduction, and transformation [7]. We next tested which mechanism was the most prevalent in the HGT events we identified in our analyses. To do this, we repeated the same analysis separately for chromosomal DNA and plasmid DNA. Our analysis reveals that the match length distributions considered in this study are strongly enriched for sequences associated with plasmids (see Fig. S7). This suggests that conjugation—mediated transfer involving a plasmid—is the primary mechanism responsible for the long exact sequence matches observed between different bacterial genera. Therefore, we should interpret the effective mobilome length in our analysis as the total length of the genes frequently located on plasmids. In the following Section we discuss the inferred effective length of the mobilome in more detail.

### C. Abundance of transferred sequences is broadly distributed, leading to short effective mobilomes

Our model assumes a mobilome of fixed size in which all elements transfer at the same rate. Fitting the MLDs yields a direct mobilome size of *L*_*d*_ ≃ 10^5^ bp and a hub-associated mobilome size of *L*_*h*_ ≃ 2.2 × 10^4^ bp, both are substantially smaller than a typical bacterial flexible genome size. To understand the origin of this unexpectedly small effective mobilome, we examined the relative abundance of horizontally transferred sequences across different genera. After clustering all identified matches into homologous groups, we quantified the relative abundance of each group. The resulting distribution (Supplemental Figures S15 and S16) is broad and closely follows a power law with exponent −3*/*2. Although a theoretical model explaining this distribution is still lacking, the observed heavy tail indicates that certain sequences are transferred far more frequently than others, effectively reducing the mobilome to a small subset of highly mobile elements.

## V. DISCUSSION AND SUMMARY

Our study shows that match length distributions (MLDs) provide a direct readout of the HGT network structure. In particular, we show that the shapes of the MLDs are well explained by a continuous process of gene flow between bacterial genera. While only pairwise HGT rates are sufficient to account for pairwise MLDs, we show that ignoring indirect HGTs would fail to successfully model the gene exchange processes occurring between bacterial genera. In contrast, our network-based approach captures emergent properties that arise from indirect HGTs via network hubs.

The exponent of the power-law, *α*, serves as a highly informative fitting parameter. Our theoretical analysis predicts that *α* should take integer values, a prediction that is well supported by empirical observations. Furthermore, *α* encapsulates information about the underlying topology of HGT network. In particular, we show that for a set of *n* interacting genera, the exponent equals *α* = *n* + 1 when gene transfers occur predominantly within the set, and increases to *α* = *n* + 2 when one or more external hubs act as transfer mediators or as shared sources of genes for the genera in the set.

A key conceptual result of our model is that each genus can be characterized by a scalar quantity we term its *transferability*, which reflects its intrinsic tendency to donate and receive genes. The direct HGT rate between any two genera is then the product of their transferabilities. The inferred transferabilities are well correlated with the average number of plasmids per genome and with the fraction of the genome shared with the other considered genera (see Figs. S2, S3, S5). This naturally explains the observed heterogeneities in gene flow, and provides a minimal parameterization for modeling large-scale exchange networks.

The model predicts the presence of a high-transferability hub within the HGT network. This hub can be interpreted in several ways: it may correspond to one or a few taxa with exceptionally high transferability that act as sources of gene flow to the genera. Alternatively—and more plausibly—it may represent the collective effect of many low-transferability organisms that mediate indirect transfers between other genera. Our framework demonstrates that such an effective hub is required to reproduce the observed MLD patterns in real genomic data and provides a quantitative measure of its influence. Although the direct identification of these hub organisms is currently limited by the incomplete coverage of existing genome databases, it may become feasible with broader and more systematic genomic sampling.

The MLD-based approach developed here is alignment-free, computationally efficient, and robust to varying degrees of sequence divergence. These properties make it particularly well-suited for analyzing diverse microbial genomes and metagenomic datasets. Unlike traditional methods that require multiple sequence alignments or phylogenetic reconstruction, our method directly leverages raw sequence similarity patterns. However, in this study we consider only the network of bacterial taxa (genera). We ignore any ecological, genetic, and other important differences between strains belonging to the same genus. Therefore, the HGT rates are effective, in the sense that they are averages over many strain-specific rates. Although we have focused on bacterial genera, the framework we propose is could be applied to other systems where horizontal transfer occurs, like microbial communities, virus-host interactions, and synthetic biology constructs.

In sum, we present a quantitative, predictive framework that connects genomic signatures to evolutionary processes on HGT networks. By linking match length statistics to network topology, our approach enables principled inference of microbial evolutionary web of life from large-scale sequencing data. This work lays the foundation for a broader understanding of genome evolution in the context of structured gene flow.

## VI. METHODS

### Data

We used seven genera from the *Enterobacteriaceae* family and one close outgroup: the *Serratia* genus. To see how the power law breaks into a mixture of exponentials, we added the genus *Cronobacter*, which has a small number (46) of sequenced genomes in the database, and the phylogenetically distant genus *Vibrio*. We used only complete genome-level assemblies in the RefSeq [38] database via NCBI [39]. To mitigate sampling biases we removed samples obtained from large multi-isolate projects, as they are indicated in the NCBI site. The list of genomes with their accession numbers and the bash script to download the data is deposited on `github` [40].

### A. Finding maximal exact matches

To find the maximal exact matches shared by genera pairs (sets with *n* = 2) we used `bfmem` software [41]. To find matches for sets with *n* > 2 we intersect the matches found for pairs. The script to do so can be found in the `github` repository [40].

### B. Calculating and fitting MLDs

For the MLDs we used logarithmic binning with 20 points per decade. Namely, our bins edges are (1000 − 0.5) · 10^0.05*k*^ where *k* is an integer. In the plots, the centers of the bins are the geometric means of their edges. Within each bin we count the number of matches and normalize it by the size of the bin. In addition, we normalize the MLD by the total number of alignments we perform for the considered set. If we analyze set *s* of size |*s*| = *n* with *k*_1_, *k*_2_, …, *k*_*n*_ genomes in the database, we divide the total MLD by *k*_1_*k*_2_ · · · *k*_*n*_.

To fit the parameters of the model we minimize the mean squared log-difference between the empirical and the predicted MLDs. The mean is 15% trimmed: to make it more robust, the worst 15% of the bins for each set are not included in the mean. The minimization was performed in two steps: global GA [42] and then L-BFGS-B optimization using the `optimParallel` package [44] in `R`.

The estimate of the power-law exponent *α* for each set is obtained using the method-of-moments estimator [45]. All relevant scripts can be found in `github` repository [40].

## Acknowledgments

We thank Eugene Koonin and Lev Melnikovsky for useful discussion and suggestions. Calculations were performed on the Weizmann Institute’s WEXAC supercomputer and cluster “Afalina” at Sevastopol State University. M.S. was supported by the European Union (ERC, BIGR, 101125981), Israeli Science Foundation (146873) and the Clore Center for Biological Physics.

## Supplemental Results

### S.1. GENERAL EQUATION AND ITS ANALYTIC SOLUTION

In this section we present the model and derive in more detail analytical results for the expected match length distribution (MLD) for a given HGT network. As network nodes we use bacterial taxa (we use genera in this article, but in this section we keep it more general), while the edges represent the effective rates of HGT between the taxa pairs. Effective rates take into account that the transferred gene is not present in 100% of the genomes of the donor taxon and is not transferred to 100% of genomes of the recipient taxon. We assume that all transfers are Poisson processes with different rates and focusing on long matches shared by distant taxa, thereby neglecting the contribution of conservation to the exact sequence matches. Throughout the paper we define time units such that the mutation rate *µ* = 1, such that all inferred effective HGT rates are given in units of a typical bacterial mutation rate *µ*.

We consider network of taxa exchanging genetic material from a mobilome of length *L*. The exchange rate from taxon *i* to taxon *j* is denoted by *ρ*_*ij*_. We obtain the analytic solution for a general asymmetric matrix, but to fit the model we assume that every taxon *i* is characterized by its transferability *γ*_*i*_, such that the transfer matrix is symmetric and rank-1, given by *ρ*_*ij*_ = *γ*_*i*_*γ*_*j*_.

To simplify the notation we define *ρ*_*ii*_ = 0. For a set of distinct taxa *s* = {*i, j, k*, … } with *i* < *j* < *k* < …, we define *P*_*s*_(*τ*)*dτ* as the expected fraction of the mobilome for which the total length of the phylogenetic tree of the set *s* is in the interval [*τ, τ* + *dτ*]. For a trivial set of a single taxon, obviously, *P*_{*i*}_(*τ*) = *δ*(*τ*) ∀*i*. For completeness we solve separately the case with the additive HGT and replacing one, although the asymptotic behaviour is the same in both cases and for the regime that is relevant for our study both scenarios are practically indistinguishable.

#### S.1.1. How to relate *P*_*s*_(*τ*) to genomic data

Before we proceed to solve the model, we discuss here how to validate our model using empirical data: how to relate calculated *P*_*s*_(*τ*) to an empirical quantity. In principle, using the molecular clock assumption, evolutionary time divergence *τ* between two DNA loci is related to the fraction of mismatches. However, as stated above, our model results in mosaic structure of the genome, such that different loci possess different time divergences. Thus, a simple molecular clock with a single divergence time cannot be used. Instead we have a combination of clocks in different loci—each with a different divergence time. Furthermore, for real bacterial genomes we don’t know where one locus ends and another one starts, so one cannot calculate *P*_*ij*_(*τ*) directly from two bacterial genomes to compare it to the one that we derive below using our model. A method to circumvent this problem was presented and used in Refs. [30, 46–49]. The main idea is to use statistical properties of the exact sequence matches between the homologous sequences of a pair instead of the statistical properties of time divergence. One can show [50], using the result derived in [35] that the molecular clock instead of fraction of mismatches can be formulated in terms of the distribution of exact matches lengths *r*. Namely, for a given divergence *τ* between two loci of length *K* the expected number of their exact sequence matches of length *r* ≪ *K* is given by

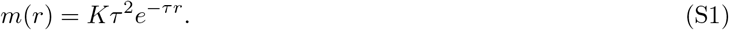

Therefore, if the divergence probability between strains *i* and *j* is *P*_*ij*_(*τ*) the exact match length distribution of *i* and *j* genomes of length *L* (with ≃ *L/K* loci) is given by

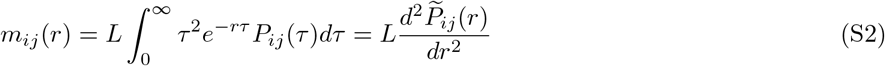

where 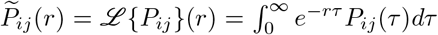 is the Laplace transform of *P*_*ij*_(*τ*). One can see that length distribution of exact sequence matches between two sequences is related to the Laplace transform of their time divergence distribution. This is why in the following we analytically calculate not only *P*_*ij*_(*τ*) but also *m*_*ij*_(*r*) because the last quantity can be relatively easily obtained from the empirical data, in contrast to *P*_*ij*_(*τ*).

In the following we split our analysis into two different types of HGT: additive and replacing [51, 52]. The asymptotic, large *r* results will be identical in both cases.

#### S.1.2. Additive HGT

We consider a set of taxa *s* exchanging genetic material from a mobilome of length *L*. To simplify the notation we define *ρ*_*ii*_ = 0 and *P*_{*i*}_(*τ*) = *δ*(*τ*) ∀*i*. So far we implicitly referred to *P*_*s*_(*τ, t*) without the time dependence *t*, i.e. we referred to the steady state of the tree length distribution. We are interested in the steady state, as it does not depend on the sampling time. Here we consider the scenario when the HGT event does not replace its homolog in the recipient but simply adds the transferred sequence to the recipient’s genome. In this case the time evolution of *P*_*s*_(*τ, t*) can be written in a recurrent form. The recurrent form is expressed in term of *P* of one of the two following sets: {*i, s\j*} sets where taxon *j* ∈ *s* from the set *s* is replaced by taxon *i* ∉ *s* (i.e. incoming gene), and the subsets of the kind *s\j* which are obtained removing taxon *j* ∈ *s* from *s*.

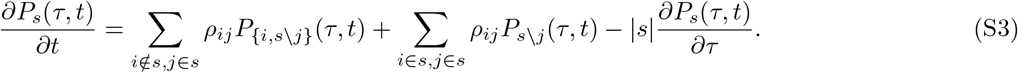

The first term represents the transfers from taxa *i* ∉ *s* outside of *s*, while the second term represents transfers within the set. The third term represents accumulation of the total evolutionary distance of the set |*s*| with *s* members. Because the HGT is additive, the sink term in the equation is absent.

Defining the matrix *R*^a^ in the sets space (superscript *a* stands for “additive”),

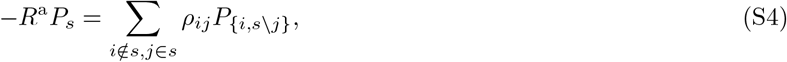

in the Laplace *r*-space, the steady state is given by

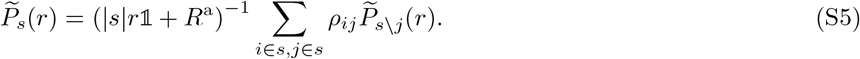

Therefore, one can calculate 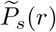 of all sets *s* with *n* members using only information about sets with *n* − 1 members. Starting from trivial sets with one member 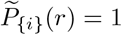 one can iteratively calculate 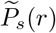 for all subsets *s* of the taxa network using Eq. (S5). Then, defining time such that the mutation rate is *µ* = 1, one can get the expected MLD of set *s* using (S2):

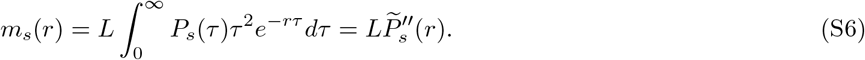

#### S.1.3. Replacing HGT

In the replacing scenario for a set of distinct taxa *s* = {*i, j, k*, …} with *i* < *j* < *k* < … the evolution of *P*_*s*_(*τ, t*) is given by

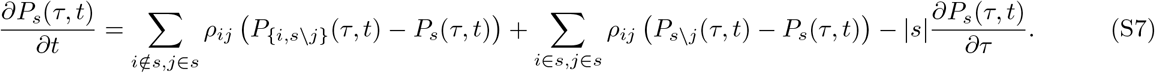

In the steady state one can write

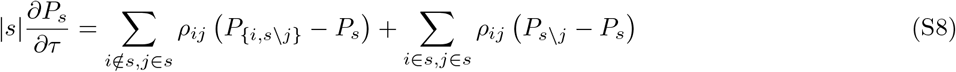

Defining the matrix *R*^r^ in the sets space (superscript r stands for “replacing”)

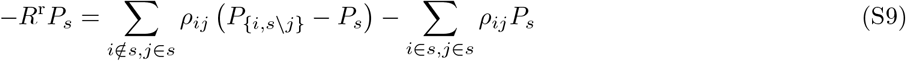

in the Laplace *r*-space

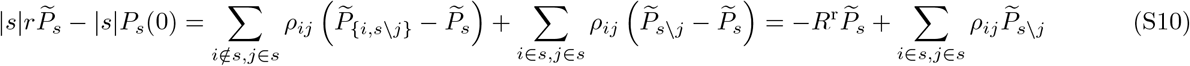

one can simplify to

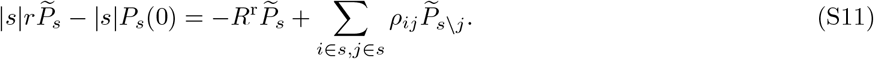

Demanding 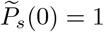 we get

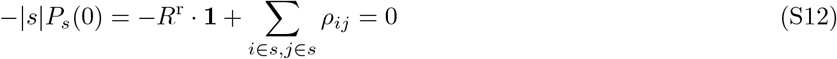

such that

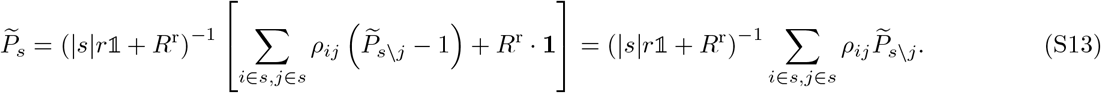

Thus, the recursive equation for 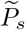 in the replacing HGT case is the same as for the additive HGT case, but *R*^r^ matrix (S9) is different from the *R*^a^ matrix for additive HGT (given in Eq. (S4)).

#### S.1.4. Asymptotic results for both additive and replacing HGT are the same

One can easily see that Eqs. (S5) and (S13) in the large *r* limit do not not depend on *R*^a^ or *R*^r^ and become identical. Below we give explicit solution in this asymptotic regime for both additive and replacing HGT scenarios, first for *s* of different sizes and later for a general set *s*.

##### S.1.4.1. *Two taxa n* = |*s*| = 2

In the limit *r* → ∞ Eqs. (S5) and (S13) for *n* = 2 are reduced to

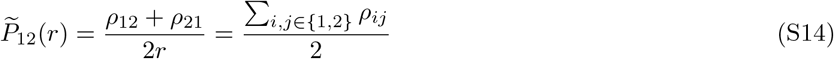

And

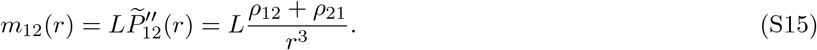

##### S.1.4.2. Three taxa

In the limit *r* → ∞ Eqs. (S5) and (S13) for *n* = 3 are reduced to

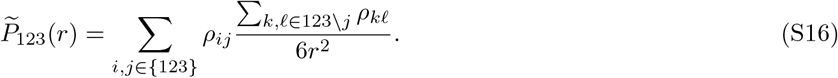

In the explicit form

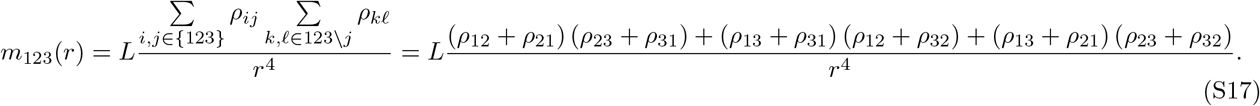

##### S.1.4.3. Four taxa

Using the same line of arguments

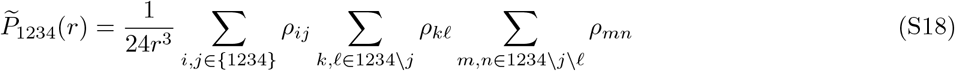

so

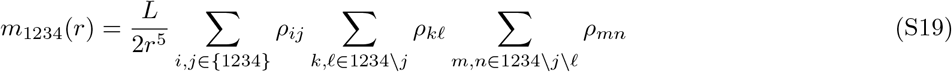

##### S.1.4.4. Five taxa

In a network of HGT with 5 taxa the MLD tail is given by

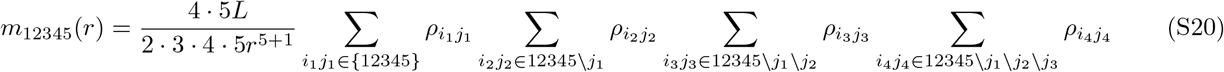

##### S.1.4.5. General asymptotic result

In a *s*-network of HGT with |*s*| = *n* taxa and asymmetric transfer rates *ρ*_*ij*_ the MLDs tails are given by Eq. (3):

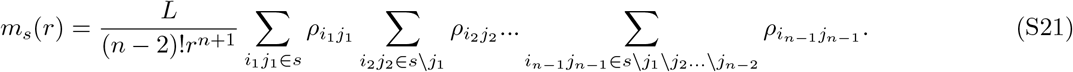

The MLD of matches shared by the same set of genera, but embedded in a larger network will have 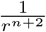 correction term to this asymptotic result. In the case when there is a strong hub outside of the set, this correction term may become dominant. In the following we calculate the 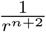 term for such a strong hub.

#### S.1.5. Common strong hub calculation

To calculate the contribution of a strong hub to the MLD we consider a hub which is connected to *n* genera. The latter have no direct connections with each other and transfer the genes only via the hub. We calculate *P*_*s*_(*τ*), *P*_*s*_(*r*), *m*(*r*) in the asymptotic *τ* →0, *r*→∞ regime. For convenience, we denote the hub with index 0 and the other genera with index *i* = 1…*n*.

Since the hub is connected directly to each genus *i* with rate *ρ*_0*i*_, in the asymptotic regime, as we found above

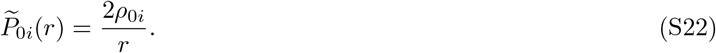

To find the pairwise MLDs we write

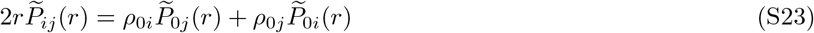

or

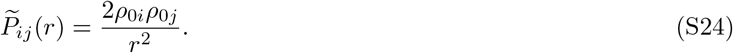

To find the MLDs of 3 genera we write

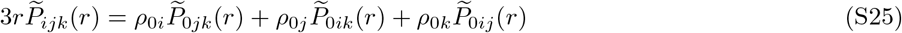

or

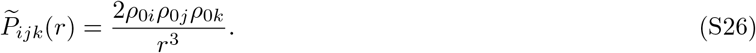

In the special case of *ρ*_*ij*_ = *γ*_*i*_*γ*_*j*_ we get

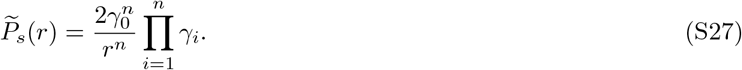

and match length distribution in the limit *r* → ∞ is given by Eq. (5)

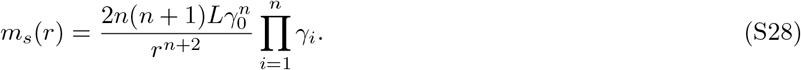

#### S.1.6. Common source calculation

In the calculation above we considered the edges between the hub and the other *n* nodes of the network as symmetric. Here we analyze the case when the HGT can occur only from the hub to an other node, but not in the opposite direction. In this case the hub acts as a common source of genes. We show now that in the asymptotic regime a common source can be viewed as a simple hub with transferability divided by 2^1*/n*^. Consider a common hub (with index *i* = 0) which act as a gene donor to *i* = 1, …, *n* genera with rates *ρ*_0*i*_ = *γ*_0_*γ*_*i*_. Let’s say that the HGT from the hub to the genera occurred times *τ*_1_, *τ*_2_, …, *τ*_*n*_ ago. For *τ*_*i*_ the indices *i* indicate not the taxa, but the rank of the transfer event, such that *τ*_1_ ≥ *τ*_2_ ≥ … ≥ *τ*_*n*_. The sum of all times at which mutations could accumulate is given by

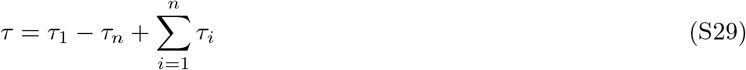

and the distribution of *τ* in the *τ* → 0 limit is given by

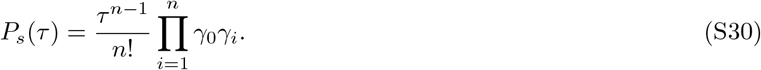

Thus its Laplace transform in the limit *r* → ∞ is

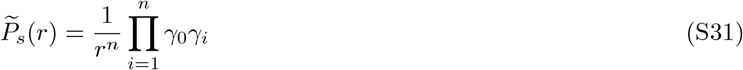

and match length distribution in the limit *r* → ∞ is given by

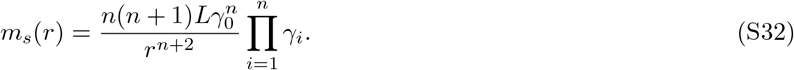

One can see that (S32) is equivalent to halved (5) or *γ*_0_ transferability divided by 2^1*/n*^. In our study we assume that the hub is connected to other genera with symmetric edges, but, as show here, our results can be easily extended to the common source case redefining *γ*_0_. The function form of the MLD is the same *m*_*s*_ ~ *r*^−(*n*+2)^ in both cases. This scaling is robust to system’s parameters and is qualitatively different from the *m*_*s*_ ~ *r*^−(*n*+1)^ scaling for the system where the transfers are predominantly within the set *s* (see Eqs. (3,4)).

**FIG. S1:**
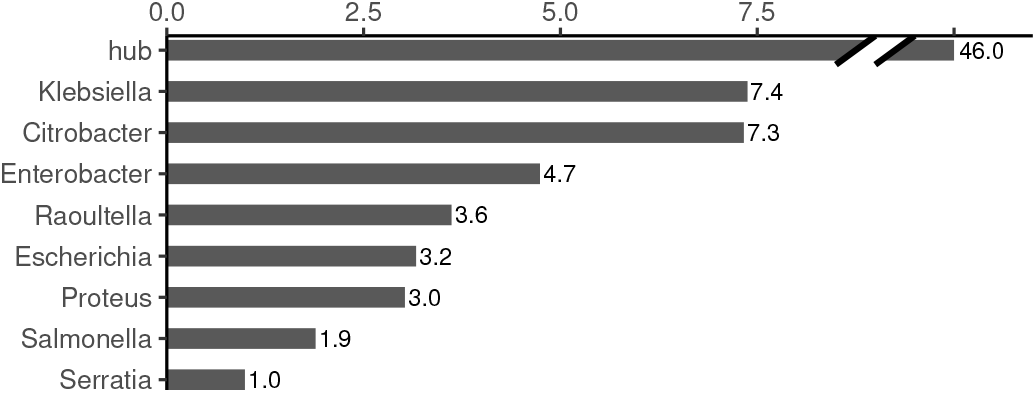
Calculated transferabilities of studied genera.

**FIG. S2:**
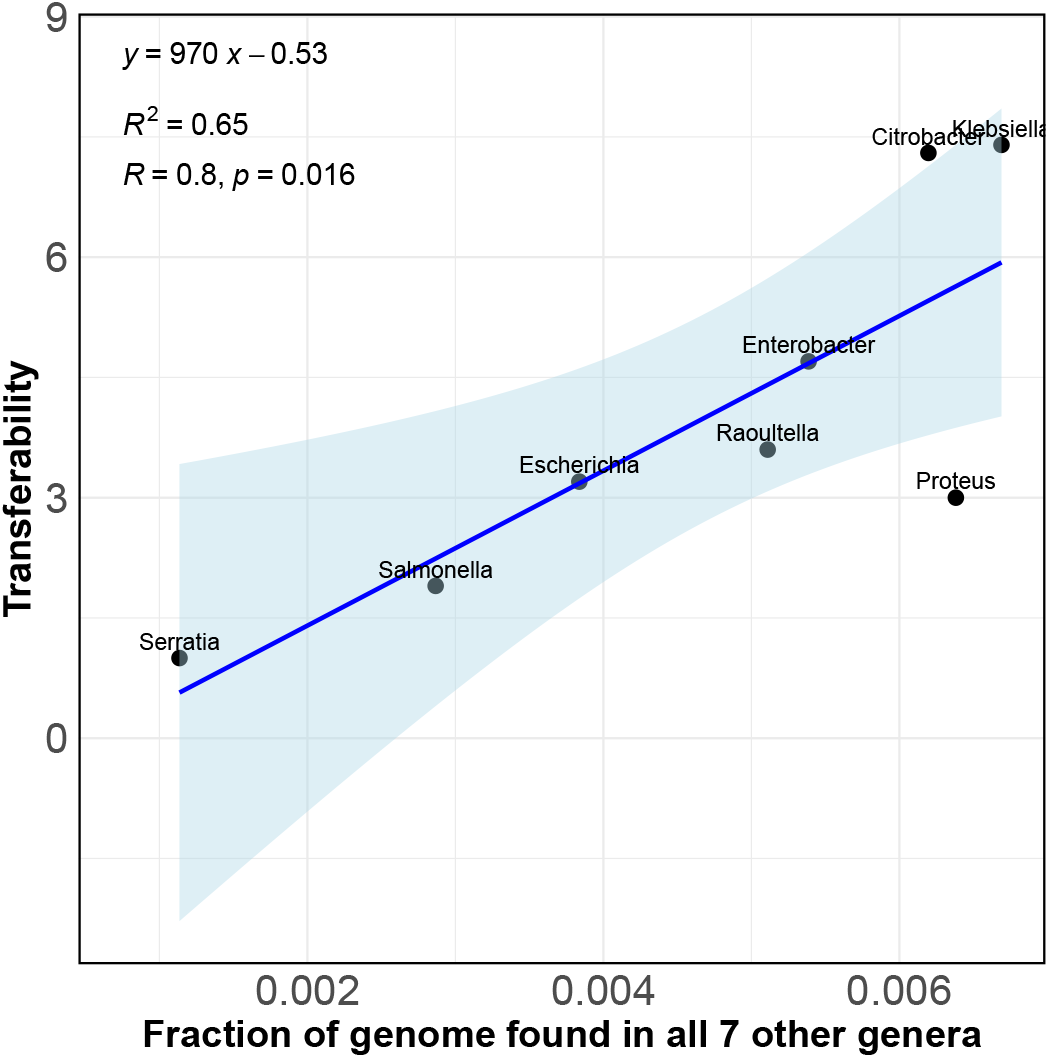
Calculated transferabilities of the eight studied genera *vs*. average fraction of genome shared with 7 other genera (at least one genome for each genus).

### S.2. MATCHES SHARED BY MANY *n* = 9, 10 GENERA

Here we analyze the matches which are shared by *n* = 9, 10 genera to test the limits of our assumptions of continuous HGT. To do so we added *Vibrio* and *Cronobacter* to the considered genera (only for this analysis) and found matches shared by genera from these augmented sets. These matches appear due to HGT events which manage to spread in significant fractions of the population in many genera. Such events are rare and, as our theory predicts, result in MLD which can be well fitted by a combination of a few exponentials.

As an instructive example consider matches shared by *n* = 9 genera: *Escherichia, Klebsiella, Salmonella, Enterobacter, Vibrio, Citrobacter, Serratia, Proteus, Raoultella*. The MLD of this comparison is shown in Fig. 4(h). One can see that the MLD can be well modelled by 3 recent events of spreading of DNA segments in the strains of considered genera. To get more insight into the details of these 3 processes we assembled separately short (1000 bp ≤ *r* < 2154 bp), medium (2154 bp ≤ *r* < 8000 bp) and long (*r* ≥ 8000 bp) matches using `metaMDBG` [55].

#### a. Long matches

Long matches are assembled to a single contig of length 19,411 bp. Functional annotation of genes along this contig (performed using `eggNOG-mapper v2` [54]) results in TraL, TraE, TraK, TrbI, and TraV: genes, associated with pili assembly and production [56]. In addition there are cytosine-specific DNA methylase, Bacterial DNA topoisomerase I and relaxase. Blasting the contig against the RefSeq database we find thousands of plasmids and bacterial chromosomes with almost a perfect match to the contig. This means that this contig is a mobile element, widespread in many bacteria species, beyond the set we are studying in this paper.

#### b. Medium matches

Medium matches assemble to 11 contigs of total length 51318 bp. The genes located at the matches are associated with fluoroquinolone, chromate, mercury and multidrug resistance, detoxification of formaldehyde, transposase, resolvase, chemotaxis and phage integrase.

#### c. Short matches

Short matches assemble to 3 contigs of total length 11575 bp. The genes located at the matches are adenine-specific DNA-methyltransferase, Beta-lactamase, chemotaxis, Cupin 2 and Transposase.

Functional analysis of matches shared by all 10 genera (*Escherichia, Klebsiella, Salmonella, Enterobacter, Citrobacter, Serratia, Proteus, Raoultella, Vibrio* and *Cronobacter*) results in MLD shown in Fig. 4(i). It is similar to MLD of *n* = 9 genera, but lacks the most recent HGT event (that contributed the longest matches). This indicate that this most recent HGT event did not manage to spread into the *Cronobacter* genus (at least to the 46 genomes we analyzed for this genus). The other two older HGT events are still visible on the MLD of all 10 genera. We find that these two HGT events contribute matches with different functional annotations, as shown in Fig. S6.

**FIG. S3:**
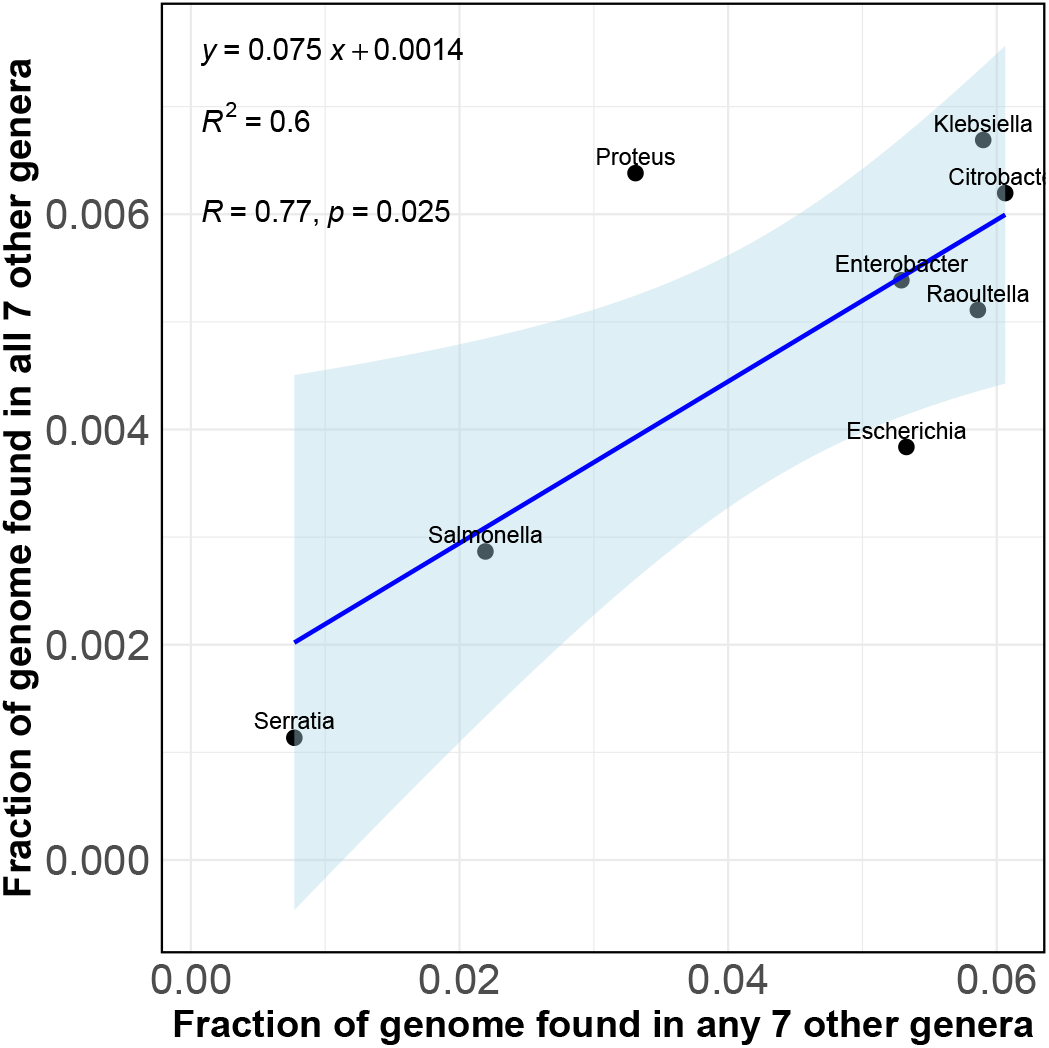
Average fraction of genome shared with all 7 other genera (at least one genome for each genus). *vs*. average fraction of genome shared with any 7 other genera (at least one genome in any genus).

**FIG. S4:**
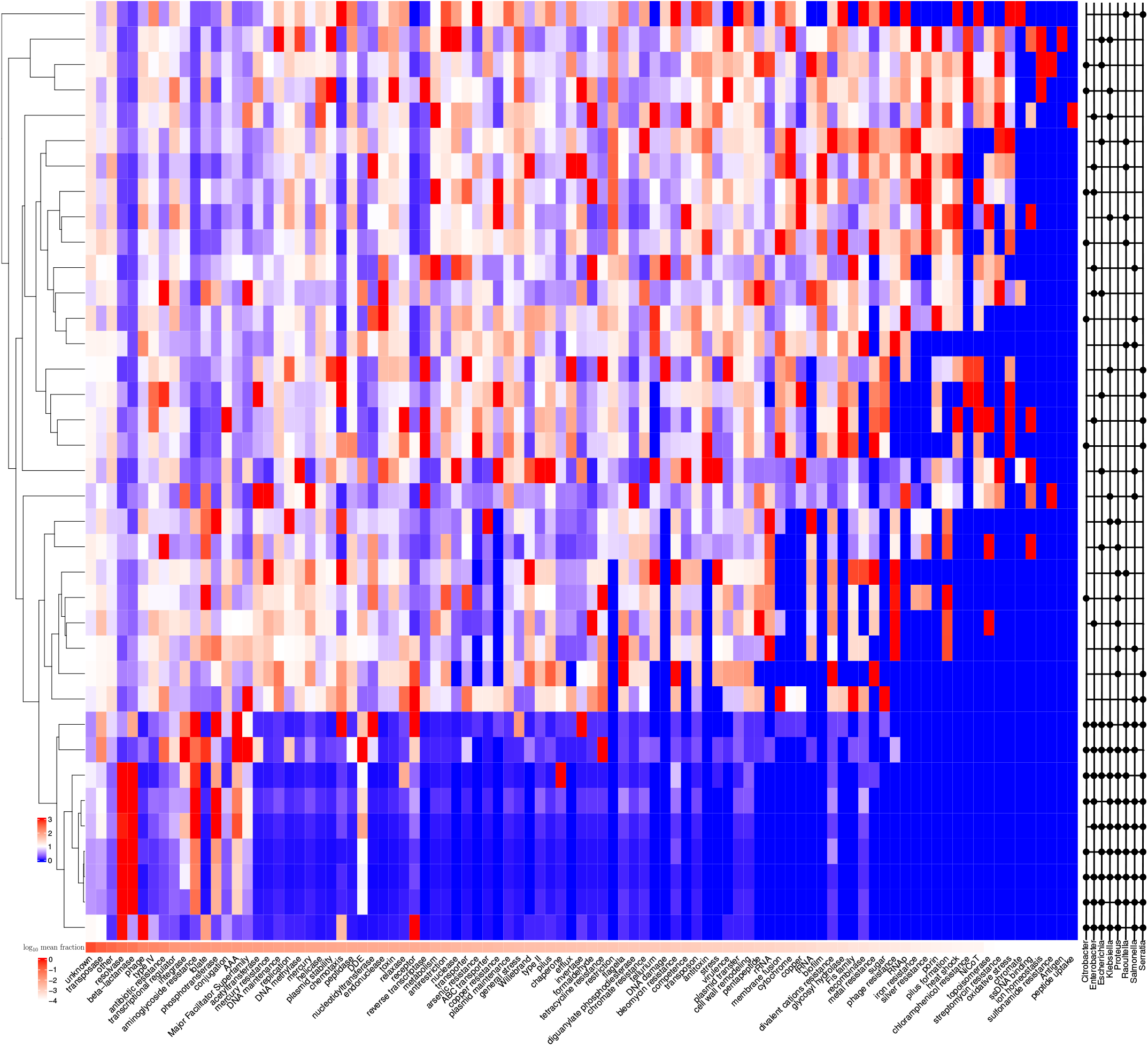
Fraction of annotations (columns) of genes located on the matches shared by sets of genera (rows). The fractions are normalized by the average fraction of all sets. To get the annotations, first for each set its match sequences were clustered using the `mmseqs easy-linclust --min-seq-id 0.5 -c 0.8 --cov-mode 1` command [53]. Resulting representative sequences were annotated using `emapper.py -i --itype metagenome -m mmseqs` command [54]. The average annotation fractions are presented in the bottom with the shades of red on the logarithmic scale.

**FIG. S5:**
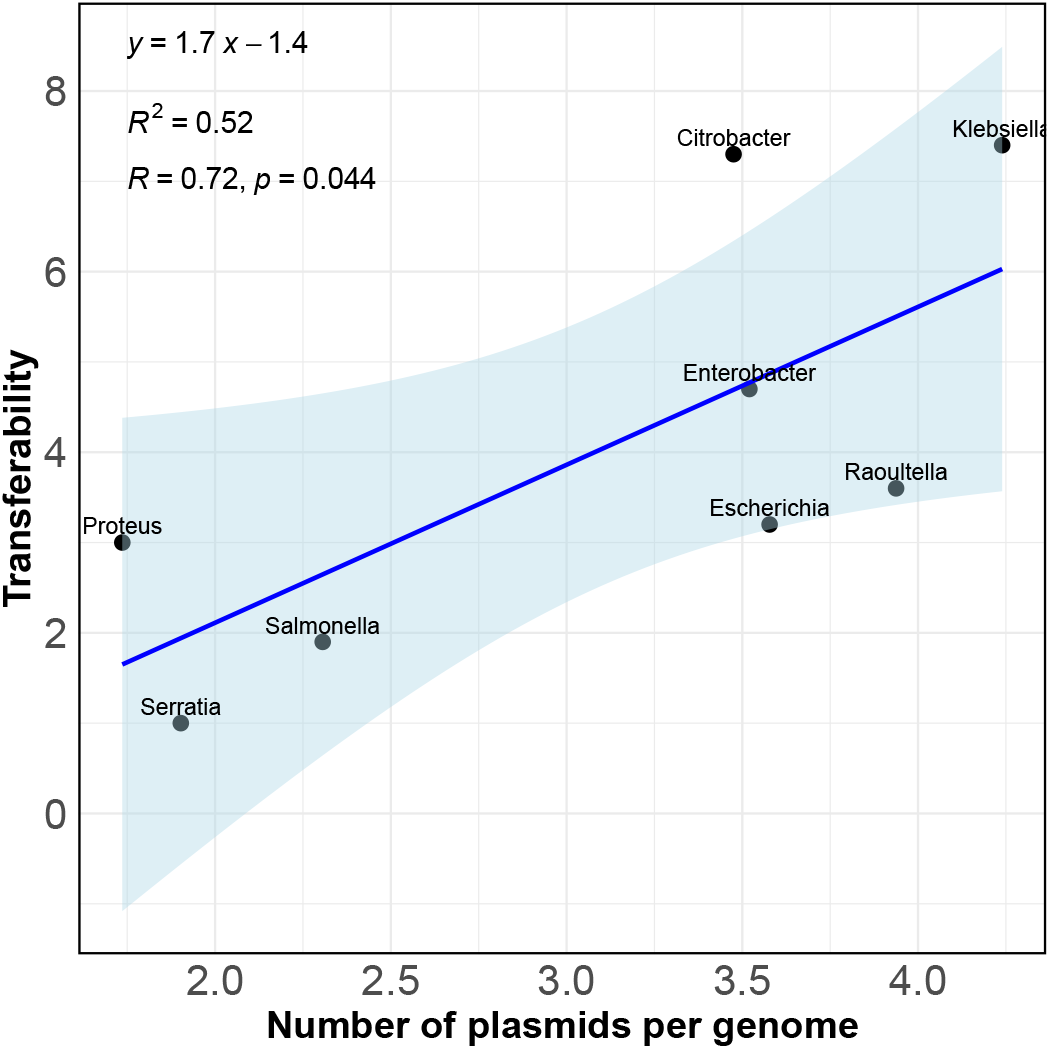
Calculated transferabilities of the eight studied genera *vs*. average number of plasmids per genome.

**FIG. S6:**
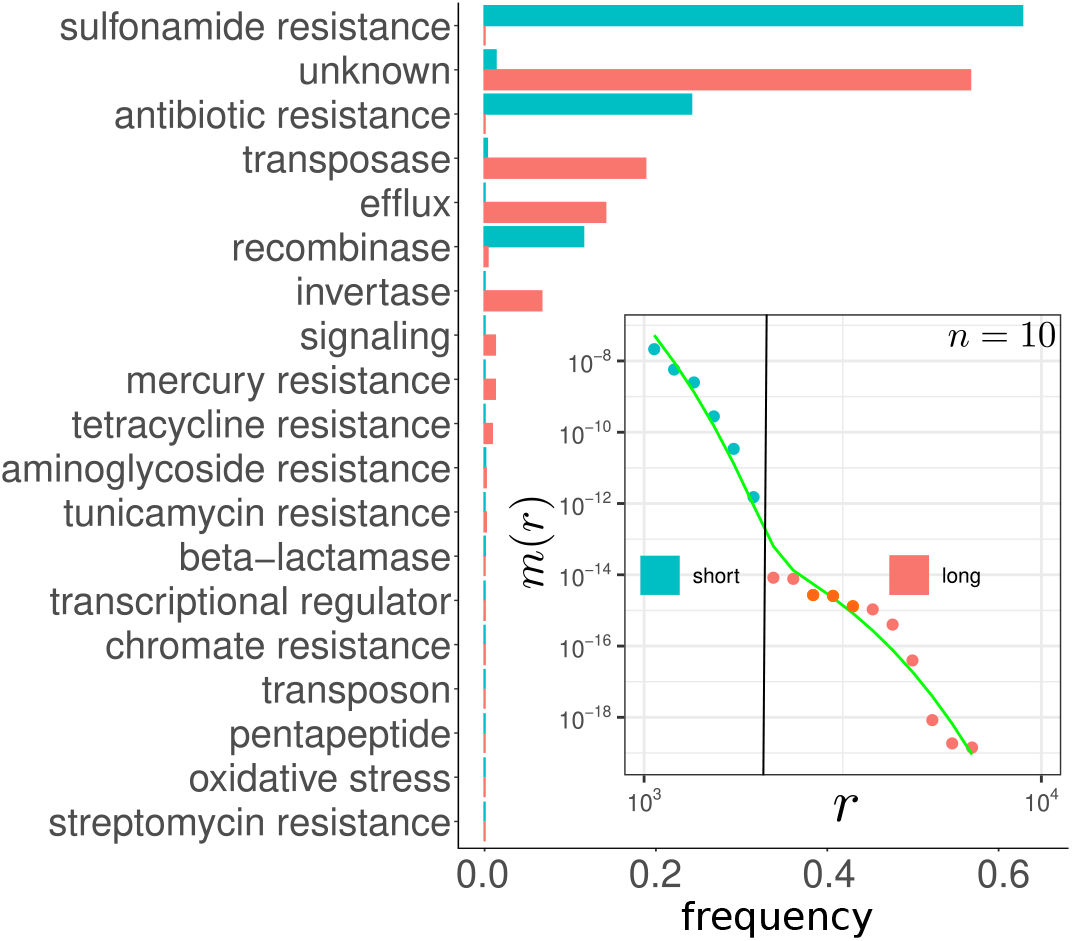
Functional analysis of matches shared by 10 genera: *Escherichia, Klebsiella, Salmonella, Enterobacter, Citrobacter, Serratia, Proteus, Raoultella, Vibrio* and *Cronobacter*. The MLD of the matches can be fitted with two exponential functions (see inset and Fig. 4(i)). Short matches (green dots and bars) and long matches (red dots and bars) have different functional annotations.

**FIG. S7:**
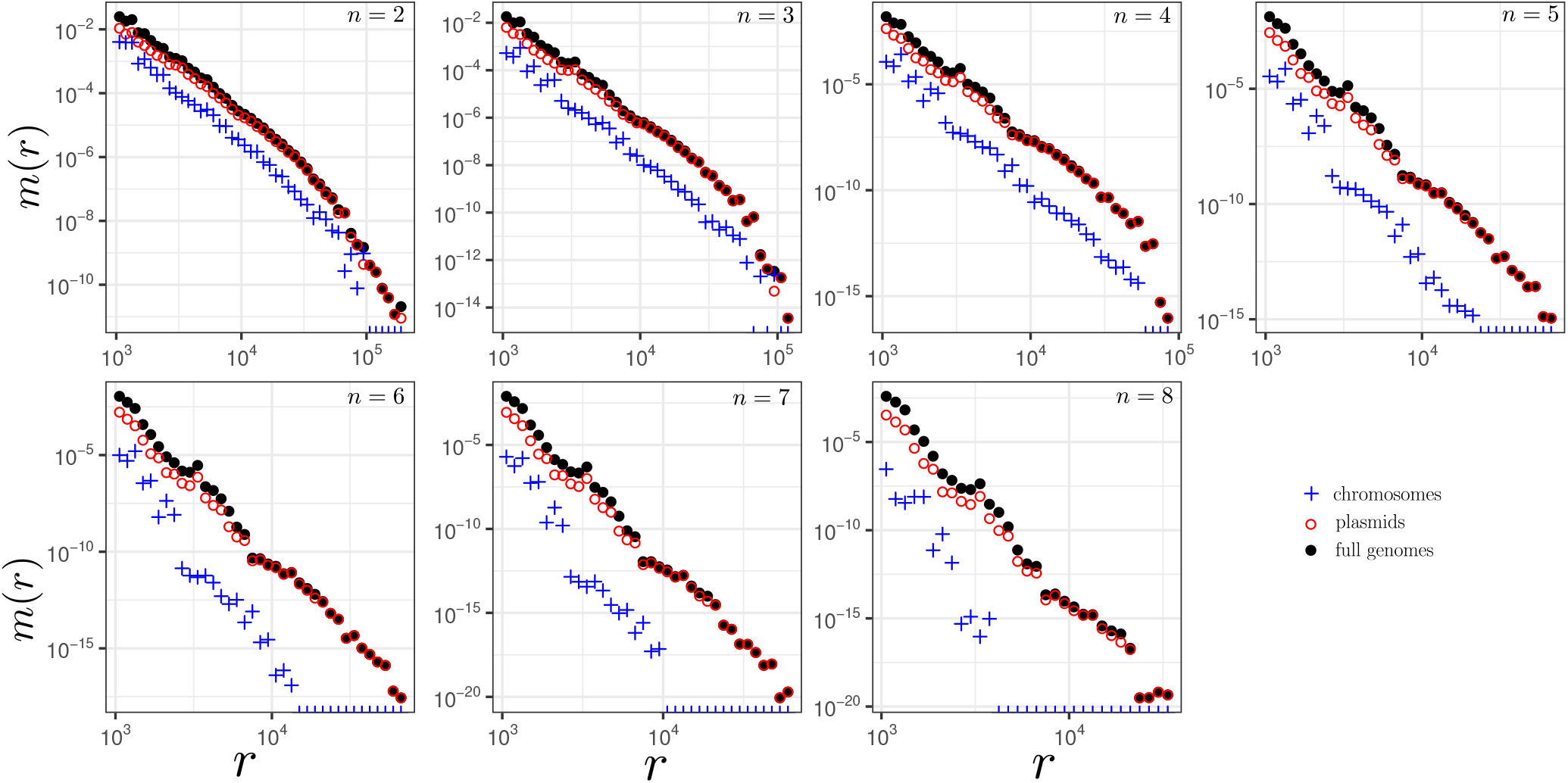
MLDs of all comparisons of pairs (*n* = 2), trios (*n* = 3), quartets (*n* = 4) *etc*. Filled black circles represent all found matches, red circles represent the matches found on the plasmids, while the crosses represent the matches along the chromosomes. Note, that crosses and red circles in these plots do not have to sum up to the black dots because the black dots also include matches between the plasmids and the chromosomes.

**FIG. S8:**
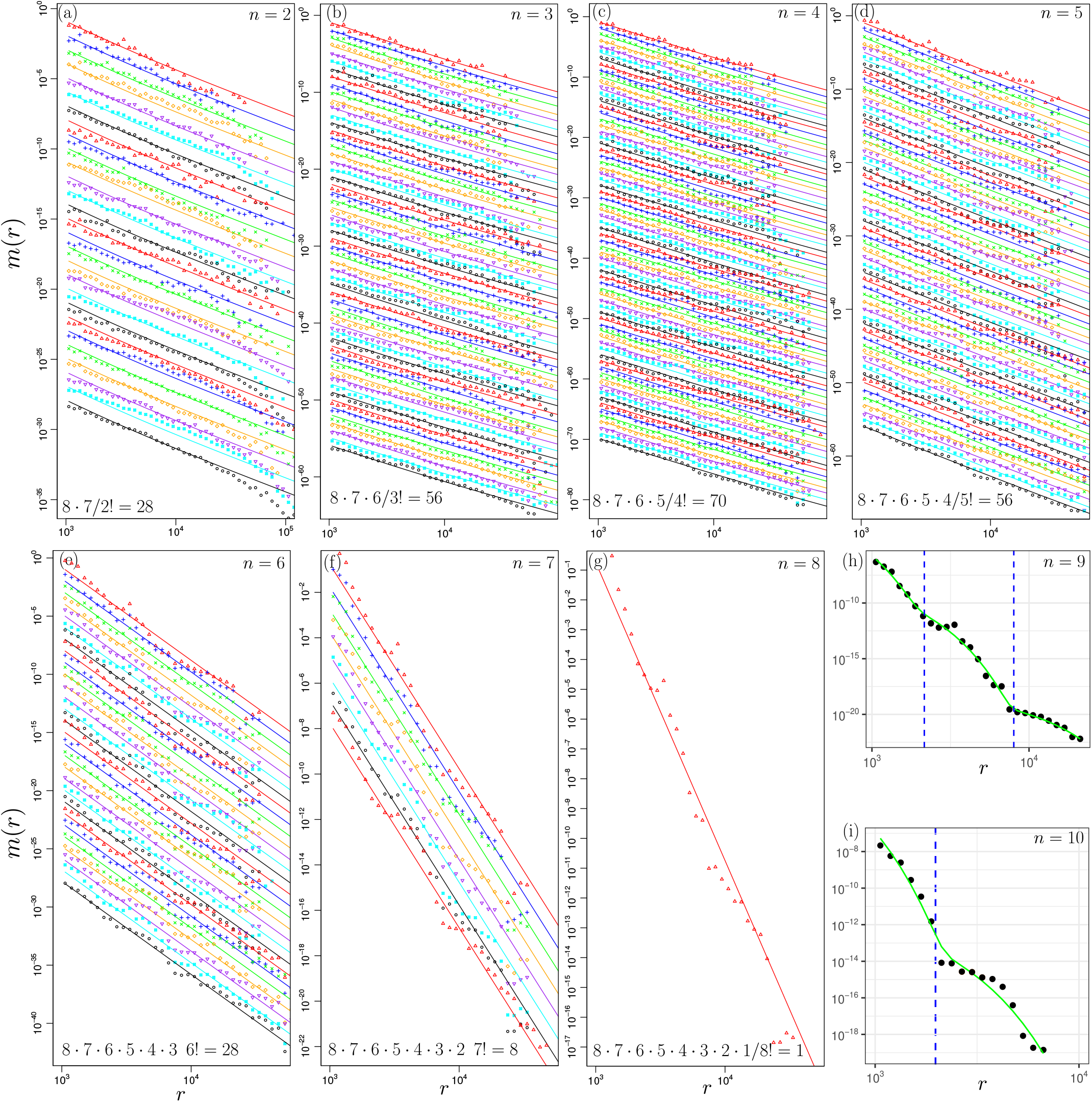
(a-g) MLDs of all comparisons of pairs (*n* = 2), trios (*n* = 3), quartets (*n* = 4) *etc* (see upper-right corners). Markers represent the empirical data and the lines represent the theoretical predictions. The MLDs are multiplies by constant prefactors for a better visibility, so the *y*-axes are arbitrary. For each set the empirical MLD and the theoretical one are multiplies by the same number. The number of considered sets for each *n* are shown in the lower-left corners. (h) MLD of *n* = 9 set: *Escherichia, Klebsiella, Salmonella, Enterobactervs. Citrobacter, Serratia, Proteus, Raoultella, Vibrio*. The green line is fit with 3 exponential functions and the dashed lines represent the crossover between the exponential functions. (i) MLD of *n* = 10 set: *Escherichia, Klebsiella, Salmonella, Enterobactervs. Citrobacter, Serratia, Proteus, Raoultella, Vibrio, Cronobacter*. The green line is fit with 2 exponential functions and the dashed line represents the crossover between the exponential functions.

**FIG. S9:**
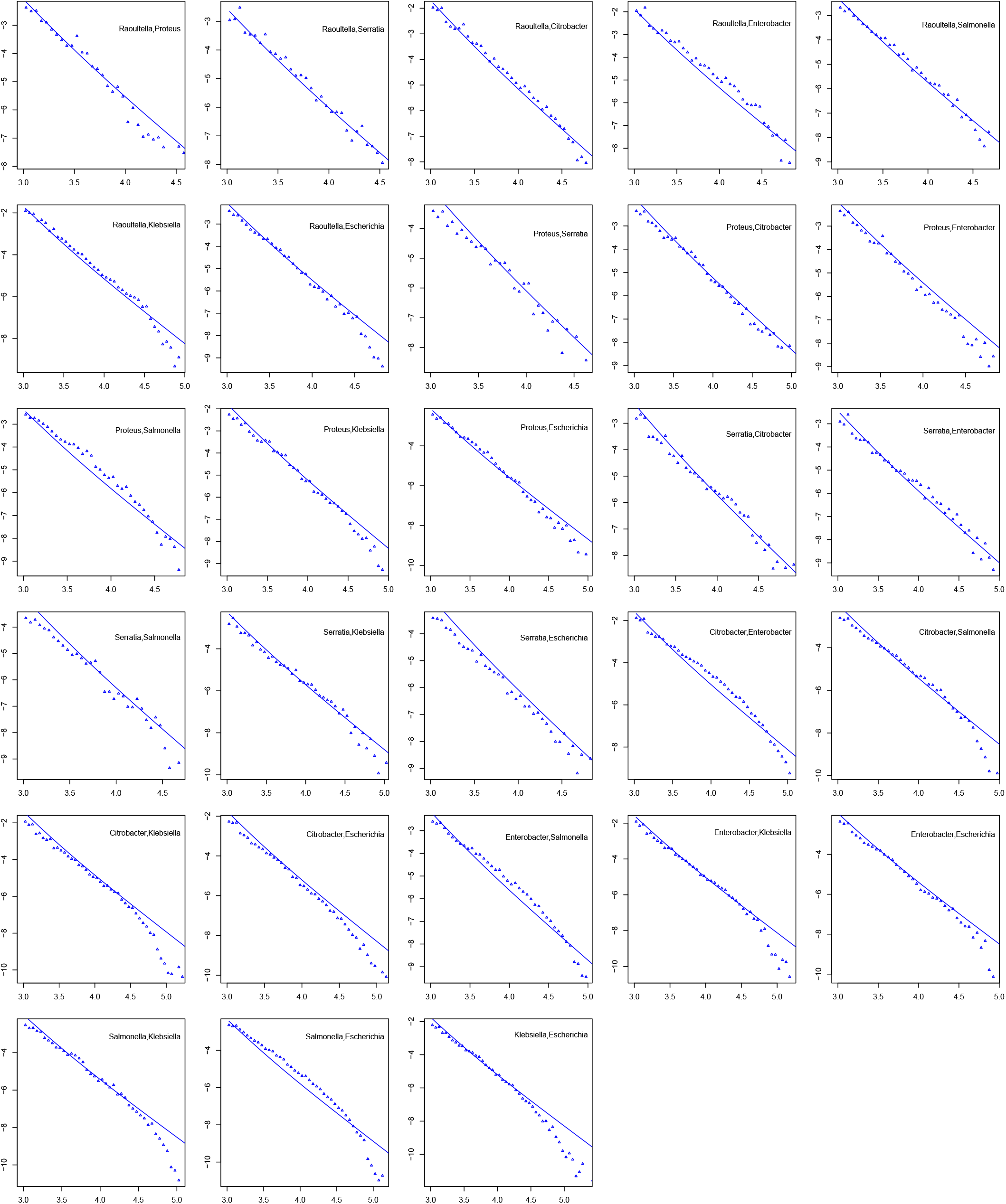
Here MLDs for *n* = 2 sets are presented on the log − log scale: log_10_ *m*(*r*) vs. log_10_ *r*. The sets are indicated in the upper right corners. Points represent the empirical data, while the lines represent the prediction of the model.

**FIG. S10:**
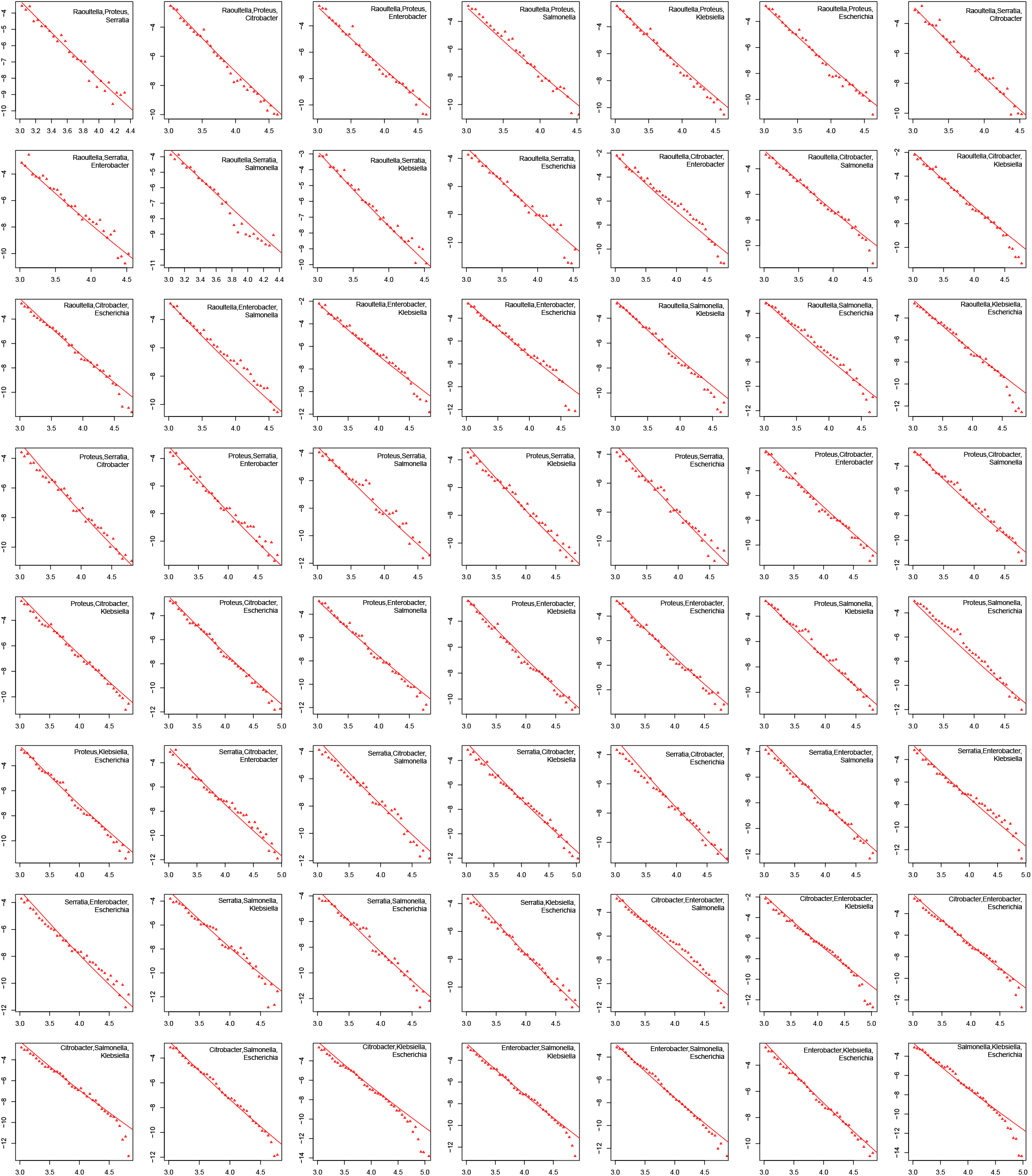
Here MLDs for *n* = 3 sets are presented on the log − log scale: log_10_ *m*(*r*) vs. log_10_ *r*. The sets are indicated in the upper right corners. Points represent the empirical data, while the lines represent the prediction of the model.

**FIG. S11:**
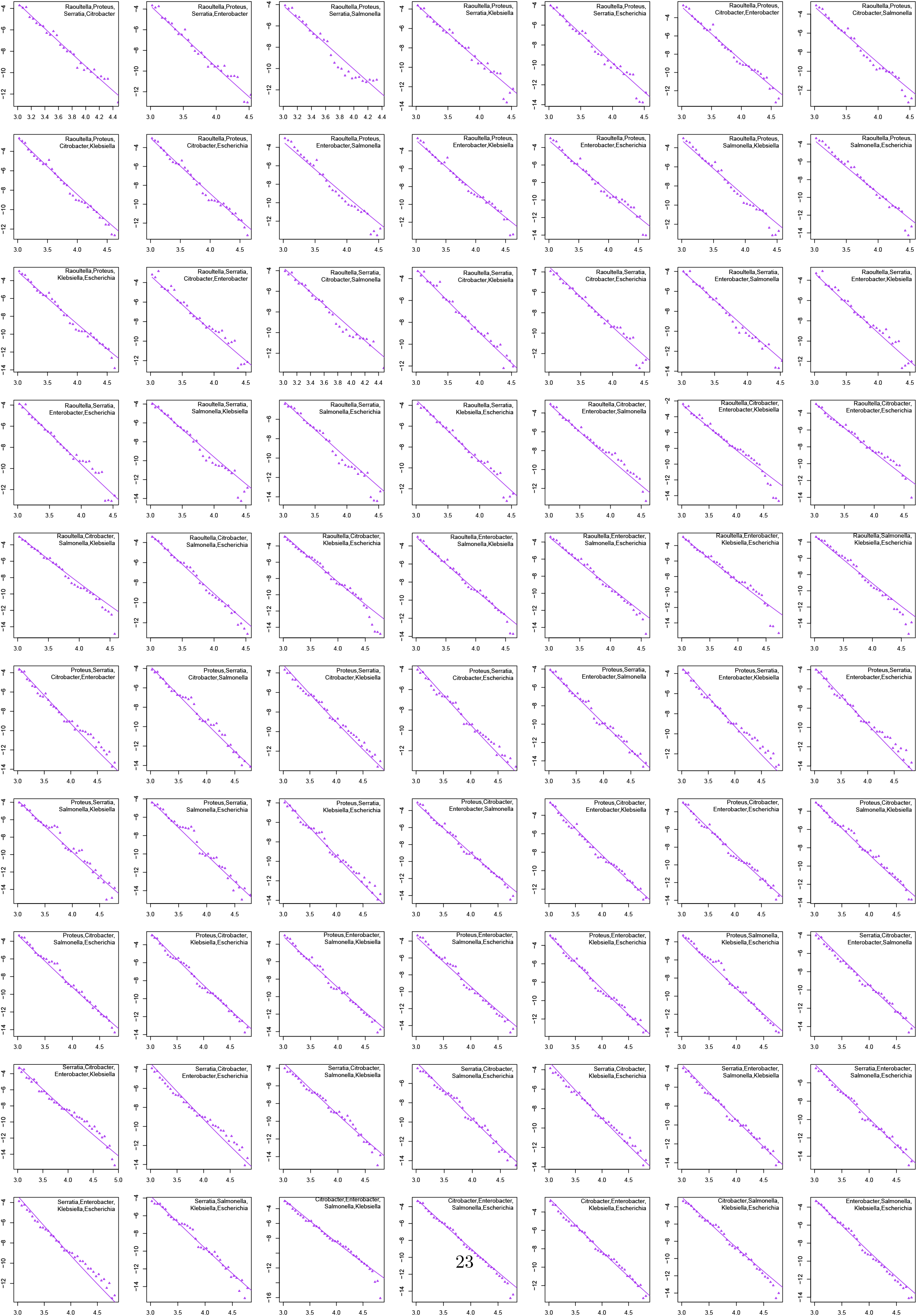
Here MLDs for *n* = 4 sets are presented on the log − log scale: log_10_ *m*(*r*) vs. log_10_ *r*. The sets are indicated in the upper right corners. Points represent the empirical data, while the lines represent the prediction of the model.

**FIG. S12:**
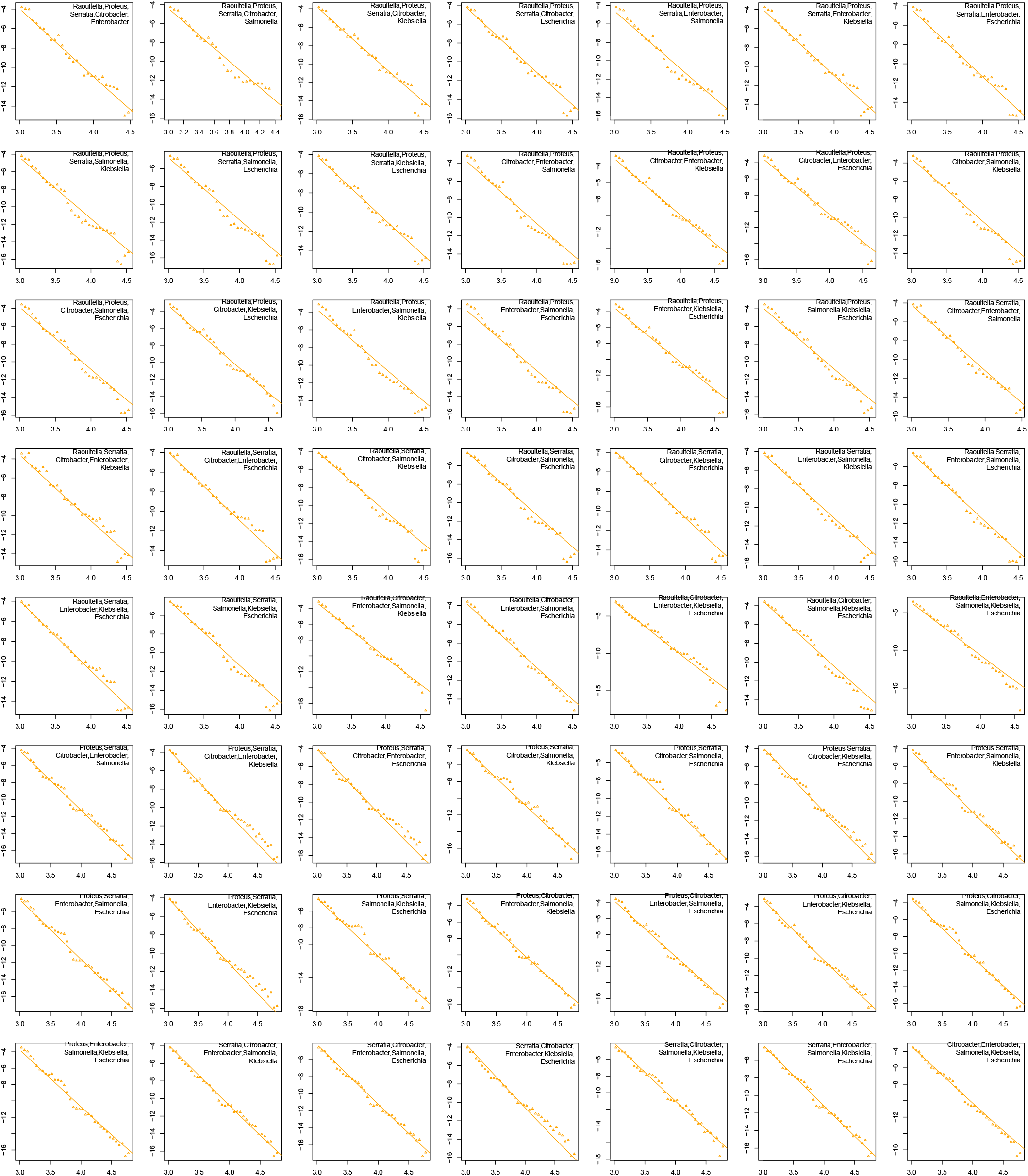
Here MLDs for *n* = 5 sets are presented on the log − log scale: log_10_ *m*(*r*) vs. log_10_ *r*. The sets are indicated in the upper right corners. Points represent the empirical data, while the lines represent the prediction of the model.

**FIG. S13:**
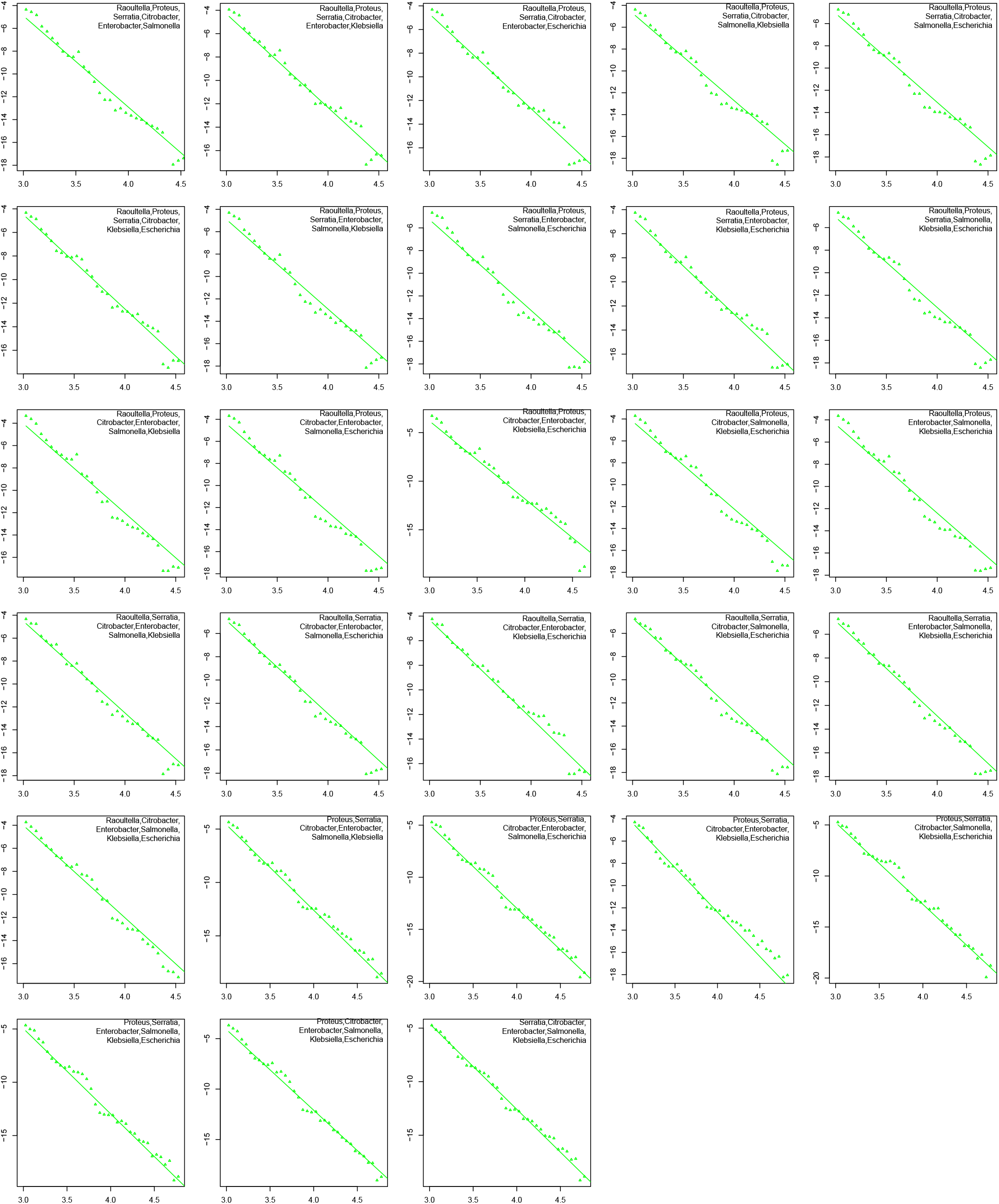
Here MLDs for *n* = 6 sets are presented on the log − log scale: log_10_ *m*(*r*) vs. log_10_ *r*. The sets are indicated in the upper right corners. Points represent the empirical data, while the lines represent the prediction of the model.

**FIG. S14:**
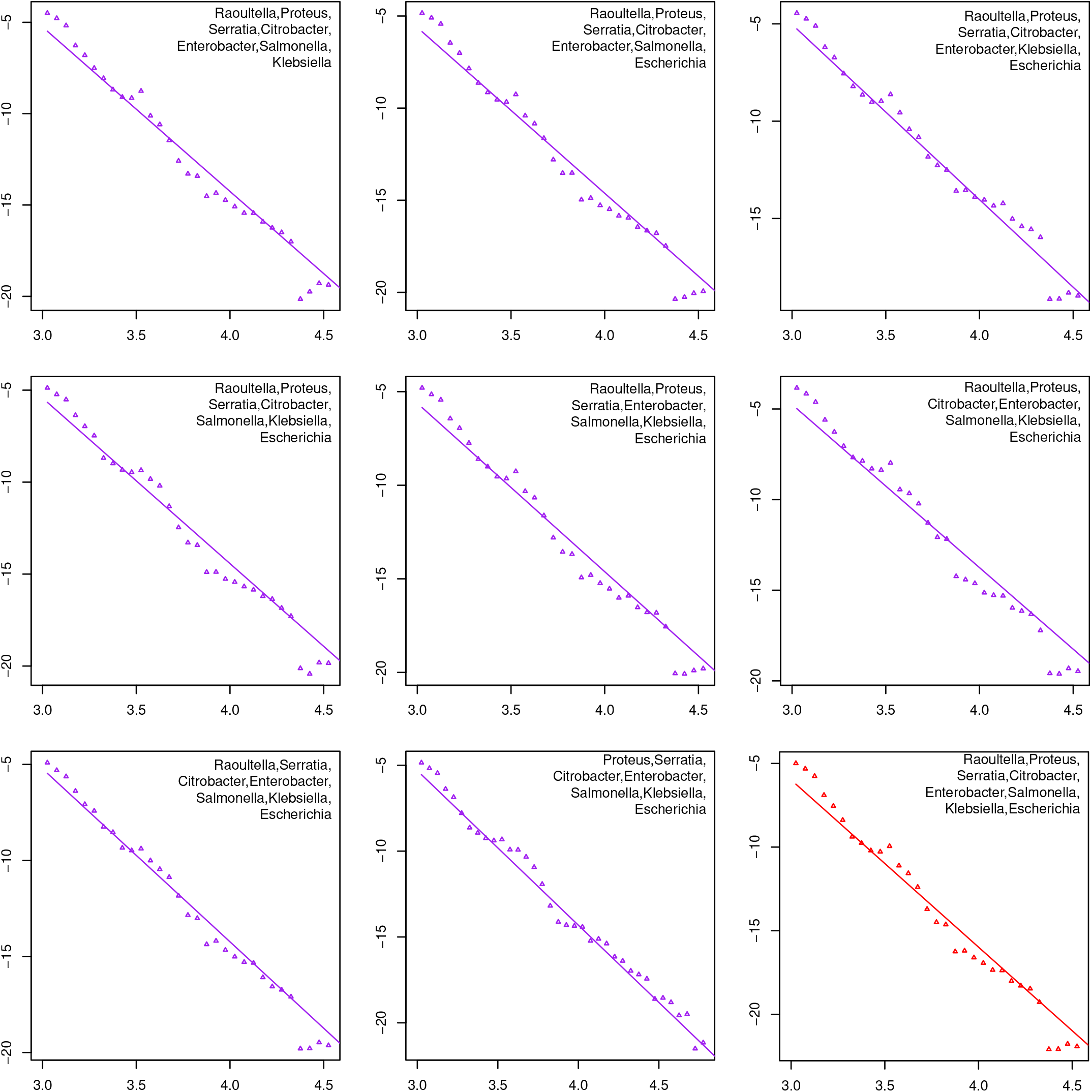
Here MLDs for *n* = 7 and *n* = 8 (last plot) sets are presented on the log − log scale: log_10_ *m*(*r*) vs. log_10_ *r*. The sets are indicated in the upper right corners. Points represent the empirical data, while the lines represent the prediction of the model.

**FIG. S15:**
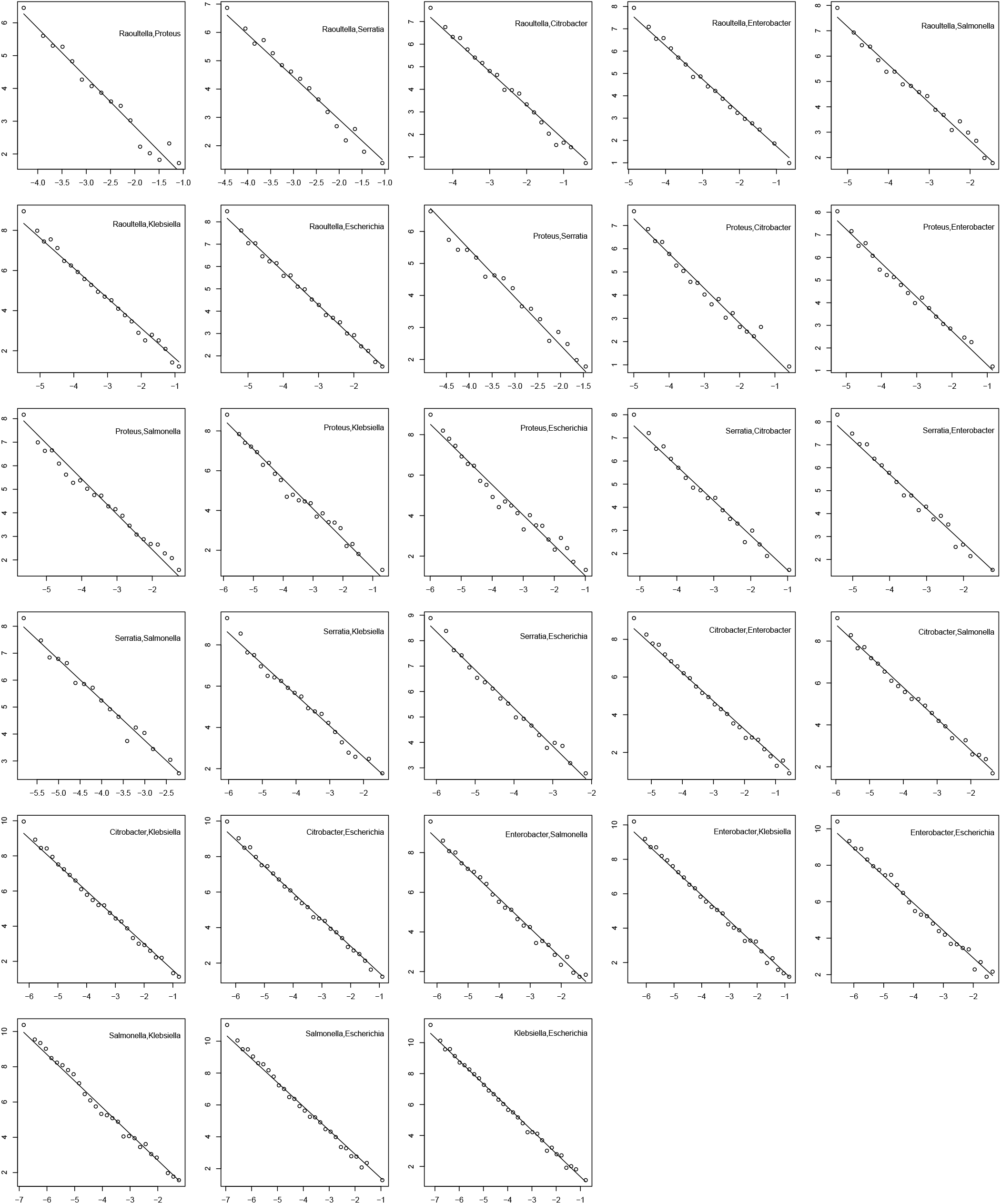
Here distributions of sequences abundances fractions are presented for *n* = 2. After clustering the matches we calculate the sizes of the clusters and divide by the total number of comparisons for each set (product of number of genomes for each genus in the set) to get the abundance fraction of each sequence—the probability to obtain this sequence taking *n* = |*s*| = 2 random genomes from the set *s*, each genome from a different genus. The probability density of this abundance fraction is presented on the plots for all pairs of genera (annotated in the upper right corners) on the log_10_ − log_10_ scale. Points represent the empirical data, while the lines represent the power-law with − 3*/*2 exponent (we don’t have a theory that predicts this exponent).

**FIG. S16:**
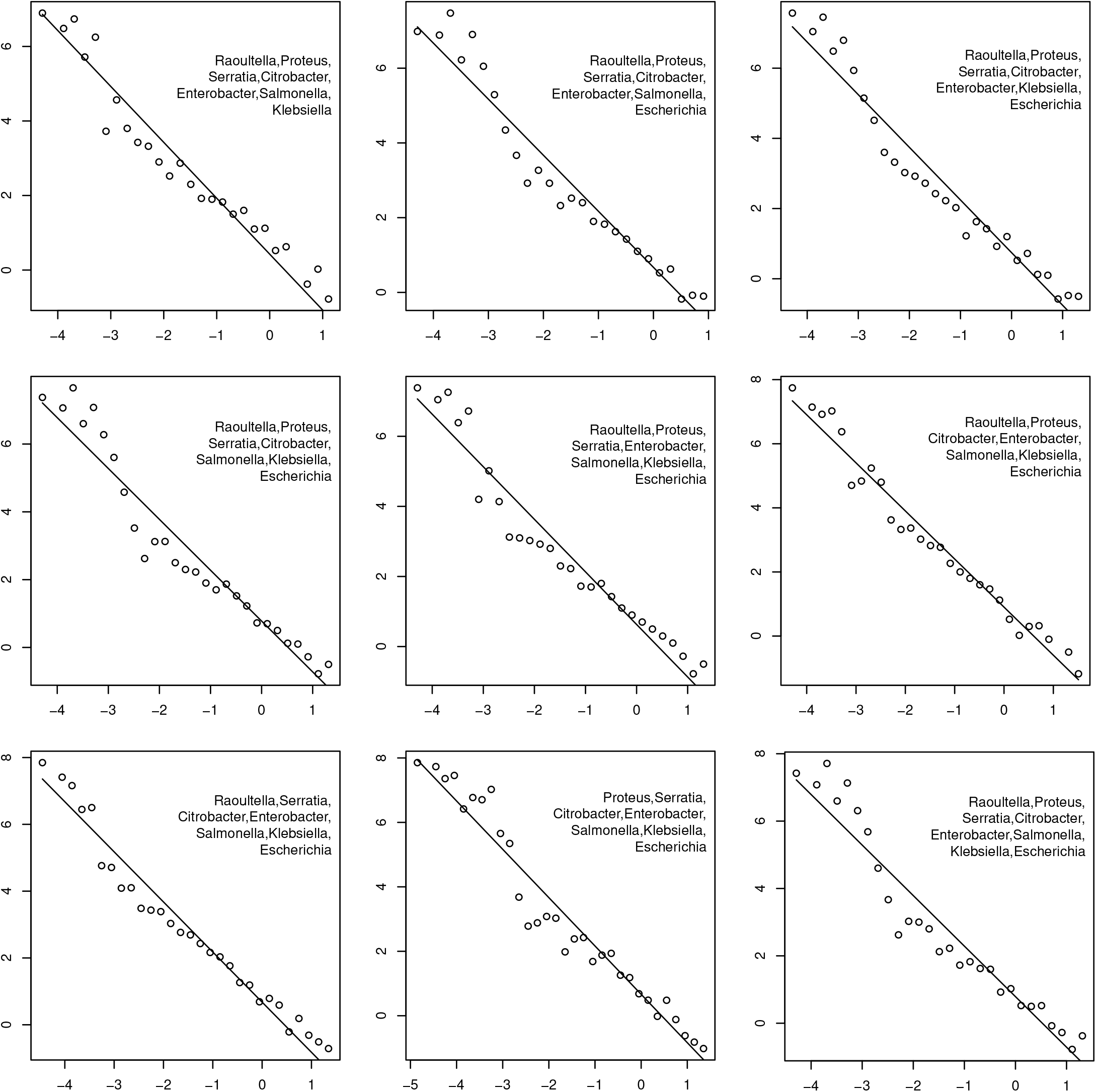
The same as in Fig. S15, but for sets of genera with *n* = |*s*| = 7 and 8 (last plot).

